# Chromosome-arm 17p Loss Renders Breast Cancer Cells Vulnerable to AURKB Inhibition

**DOI:** 10.64898/2025.12.01.691489

**Authors:** Tom Winkler, Eran Sdeor, Ron Saad, Hajime Okada, Kathrin Laue, Gil Leor, Guy Wolf, Shlomit Strulov Shachar, Uri Ben-David

## Abstract

Loss of chromosome-arm 17p (Del17p) is a genetic hallmark of breast cancer. While *TP53* loss is an established driver of Del17p, the potential therapeutically-relevant cellular vulnerabilities of this common aneuploidy remain unexplored. Here, we first analyzed genomic and clinical data from breast cancer patients using METABRIC and TCGA datasets. Del17p was prevalent across molecular subtypes and correlated with higher tumor grade, advanced stage, and worse survival. Gene expression profiling revealed reduced expression and activity of the chromosome 17p-residing gene Aurora Kinase B (AURKB) in Del17p tumors and cell lines. Moreover, functional dependency screens across breast cancer cell lines identified increased sensitivity of Del17p cells to genetic inhibition of AURKB, which we validated using chemical inhibition in matched breast cancer cell lines. Next, we generated an isogenic model of CAL51 breast cancer cells with/without heterozygous AURKB loss in *TP53*-WT and *TP53*-null backgrounds, and confirmed that heterozygous loss of AURKB resulted in its reduced expression and in increased sensitivity to the AURKB inhibitor barasertib. Notably, p53 inactivation increased AURKB expression and reduced drug sensitivity, as previously reported, but AURKB heterozygous knockout reverted these phenotypes, revealing opposite effects of common modes of p53 inactivation (Del17p vs. point mutations). Mechanistically, the phenotypes associated with barasertib treatment – mitotic aberrations, cytokinesis failure, whole-genome doubling and apoptosis – were exacerbated in Del17p cells. Our findings therefore suggest Del17p as a potential biomarker for identifying breast cancer patients who may benefit from AURKB inhibition and highlight its potential as a therapeutic target in Del17p breast tumors.

**Significance:** Breast cancer tumors with loss of chromosome-arm 17p (Del17p) exhibit reduced AURKB expression and increased sensitivity to AURKB inhibition, suggesting Del17p as a biomarker for AURKB-targeted therapy.

## Introduction

Chromosome-arm level alterations are common in breast cancer, with recurrent losses and gains affecting tumor progression, treatment response, and patient prognosis (**1-4**). Among these, Del17p stands out as one of the most frequent chromosome-arm losses in breast cancer (**2**). Del17p encompasses the *TP53* gene, a well-known tumor suppressor whose loss disrupts key cellular processes including DNA damage response, cell cycle regulation, and apoptosis (**5**). *TP53* has been established as a driver gene of Del17p in several cancer types, including in breast cancer, and *TP53* mutations and Del17p often co-occur and are considered major drivers of breast tumorigenesis (**6-8**).

Beyond the driver genes of recurrent aneuploidies, whose identification may be challenging (**9**), other cellular consequences of aneuploidy may also be harnessed for therapeutic purposes. These include either general consequences of aneuploidy, such as cell cycle dysregulation, impaired DNA damage repair, metabolic reprogramming, and dosage compensation mechanisms; or specific consequences of recurrent aneuploidies (**10**). A loss of a chromosome-arm does not necessarily result in reduced expression of genes residing on that arm, due to gene dosage compensation (**11-12**). However, in some cases, the loss of one allele does result in reduced expression of affected genes, with downstream functional consequences (**13**).

Thus, aside from the driver genes underlying recurrent chromosomal losses, co-deleted ‘passenger’ genes may also create unique cellular vulnerabilities that can be therapeutically exploited for cancer therapy (**9**). Examples of such vulnerabilities have emerged from integrative analyses of copy number alterations and pharmacological screens, leading to the identification of CYCLOPS genes (Copy-number alterations Yielding Cancer Liabilities Owing to Partial losS) (**14-15**). One such gene is *POLR2A*, which is co-deleted with *TP53* upon Del17p, resulting in increased vulnerability of colorectal and breast cancer cells to its inhibition (**16-17**). However, POLR2A is currently not a therapeutically-relevant target, and a systematic analysis and validation of CYCLOPS genes in the context of Del17p is still lacking.

AURKB is a core component of the chromosomal passenger complex (CPC) (**18**) and plays essential roles in chromosome condensation and the spindle assembly checkpoint (SAC) (**19**). *AURKB* is known to be overexpressed in many cancer types and has therefore been proposed as a therapeutic target (**20**). Several AURKB inhibitors have entered clinical trials, although none has so far reached the market (**21**). Importantly, inactivation of p53 has been proposed to be associated with resistance to AURKB inhibitors (**22-23**). Here, we identified Aurora Kinase B (*AURKB*), located on chromosome-arm 17p approximately 0.5Mb downstream of *TP53*, as a Del17p-associated vulnerability in breast cancer. We report that *AURKB* hemizygosity, resulting from Del17p, increases sensitivity to AURKB inhibition (AURKBi), and reveal that the mode of *TP53* perturbation (point mutation vs. chromosome-arm loss) is critical for this drug response. Our results suggest that Del17p status may serve as a therapeutically-relevant biomarker for identifying breast cancer patients who are more likely to benefit from AURKB-targeted therapies.

## Materials and Methods

### TCGA and METABRIC data analyses

Tumor data from TCGA (**24**) and METABRIC (**1**) were downloaded from cBioPortal (**25**). Tumor *TP*53 mutation status and Chr17p copy number were obtained from Laue and colleagues (**8**). Tumors were classified as *TP53*-mutant if at least one allele of the *TP53* locus was perturbed by any somatic alteration, including any point mutations, focal deletions, or copy number losses. Kaplan-Meier plots were generated using the R packages survival, survminer, and ggplot2. Differential gene expression analysis was performed using the DESeq2 R package (for TCGA) and the limma R package (for METABRIC). Pre-ranked GSEA was performed on the differential gene expression results using the clusterProfiler R package with the ‘Hallmark’, ‘Reactome’ and ‘Oncogenic signatures’ gene set collections from MSigDB. All other clinical comparisons were plotted using GraphPad Prism.

### Dependency Map data analysis

The aneuploidy scores (AS) of the cancer cell lines were obtained from Zerbib and colleagues (**26**). mRNA gene expression values, protein expression values, mutation status, CRISPR and RNAi dependency scores (Chronos and DEMETER2 scores, respectively) were obtained from DepMap 20Q3 release. The doubling times of the cell lines were obtained from Tsherniak and colleagues (**27**). The chromosome-arm 17p copy number status was determined for 65 breast cancer cell lines based on their copy-number segmentation data (seg. files), classifying a cell line as ‘Del17p’ if more than 80% of the chromosome-arm was lost. Cell lines were stratified into four groups based on their *TP53* mutation status and Chr17p copy number status. The DepMap portal’s two-class comparison tool was used to perform pairwise comparisons between the following groups: Del17p;*TP53*-WT vs. WT17p;*TP53*-WT, Del17p;*TP53*-WT vs. WT17p;*TP53*-mut, Del17p;*TP53*-mut vs. WT17p;*TP53*-WT, and WT17p;*TP53*-mut vs. WT17p;*TP53*-WT. Comparisons were conducted across the following datasets: gene expression, protein expression, RNAi (Broad), RNAi (Novartis), RNAi (Combined), CRISPR (Avana), and CRISPR (Sanger). For all comparisons except WT17p;*TP53*-mut vs. WT17p;*TP53*-WT, we selected the top 1% of upregulated and downregulated genes (expression) and the top 1% most sensitive genes (dependency), based on effect size (fold change). For the WT17p;*TP53*-mut vs. WT17p;*TP53*-WT comparison, we used a more lenient 5% threshold. From each pairwise group comparison, genes were retained only if they met their respective threshold (1% or 5%) in two independent dataset types (e.g., expression and CRISPR), rather than appearing only within the same modality (e.g., two RNAi screens). Genes meeting this criterion were then evaluated to identify those consistently found in the comparisons: Del17p;*TP53*-WT vs. WT17p;*TP53*-WT, Del17p;*TP53*-WT vs. WT17p;*TP53*-mut, and Del17p;*TP53*-mut vs. WT17p;*TP53*-WT, but not in the WT17p;*TP53*-mut vs. WT17p;*TP53*-WT comparison. This filtering aimed to identify dependencies specifically associated with Del17p, independent of *TP53* loss of function. When analyzing AURKB expression and dependency data, additional covariates were controlled for using the R function ‘Partialize’, including aneuploidy score, *TP53* mutation status, 17q copy number status, doubling time, and molecular subtype.

### Cell Culture

Human breast cancer CAL51 cells were cultured in DMEM (Life Technologies) supplemented with 10% FBS (Sigma-Aldrich), 1% sodium pyruvate, 4 mmol/L glutamine, and 1% penicillin–streptomycin. MCF7, MDA-MB-468, and MDA-MB-231 cells were cultured in RPMI-1640 (Life Technologies) supplemented with 10% FBS (Sigma-Aldrich), 4 mmol/L glutamine, and 1% penicillin–streptomycin. Cells were maintained at 37°C in a humidified incubator with 5% CO₂ and were cultured for a maximum of 3 weeks. Cell lines were routinely tested and confirmed to be free of Mycoplasma contamination.

### Generation of genetically engineered CAL51 models

CAL51 cells were transfected with either a pool of two guides targeting intron 1 and intron 2 of *AURKB* to remove exon 2 (which contains the translation initiation site), or with *LentiCRISPRv2 lacZ*-control and empty *LentiGuide*. Following antibiotic selection, we PCR-amplified the region flanking the two *AURKB* guides and detected a fusion band indicating successful exon removal in a subset of cells. The fusion band was of the expected size and was confirmed by Sanger sequencing to represent the fusion of introns 1 and 2. Both the *lacZ*-control cells and the *AURKB*-targeted cells were single-cell sorted into 96-well plates containing conditioned medium. Approximately 10 colonies from the *lacZ*-control single-cell sort were pooled to generate the *lacZ*-control population. For the heterozygous *AURKB*-KO cells, we assessed the presence of the fusion band in single-cell clones by PCR and Sanger sequencing and pooled 3 clones with confirmed fusion bands. Sanger sequencing of the WT band from these clones confirmed a heterozygous genotype. The WT and heterozygous *AURKB*-KO models were subsequently transfected with two *TP53*-targeting guides to knock out *TP53*. p53 KO was confirmed by RT-qPCR analysis and WB of p53 and p21. Details of all plasmids are available on Supplementary Table S1.

### RT-qPCR

Cells were harvested using Bio-TRI (Bio-Lab), and RNA was extracted according to the manufacturer’s protocol. cDNA was synthesized using the GoScript Reverse Transcription System (Promega), following the manufacturer’s instructions. RT-qPCR was performed using SYBR Green, and quantification was carried out using the ΔCT method. Details of all RT-qPCR primers are available in Supplementary Table S1.

### Western blot

Cells were lysed in RIPA lysis buffer (1% NP-40, 0.5% sodium deoxycholate, 0.1% SDS, 150 mM NaCl, and 50 mM Tris-HCl, pH 8.0) supplemented with protease inhibitor cocktail (Sigma-Aldrich, #P8340) and phosphatase inhibitor cocktail (Sigma-Aldrich, #P0044). Protein lysates were centrifuged at maximum speed for 10 minutes, and protein concentrations were measured using the Bradford assay. All samples were normalized to the same protein concentration and mixed with a sample buffer containing DTT. Proteins were separated on 10% SDS-PAGE gels. Bands were detected by chemiluminescence (Millipore, #WBLUR0500) using a Fusion FX gel documentation system (Vilber). Quantification was performed using ImageJ software or UVITEC Authentic Imaging Specialists software. Details of all antibodies are available in Supplementary Table S1.

### Immunofluorescence

Cells were seeded on coverslips in a 24-well plate, washed with PBS, and fixed for 12 min at room temperature (RT) with 4% paraformaldehyde, followed by permeabilization with 0.5% Triton X-100 for 10 min at RT. Slides were incubated with a primary antibody against AURKB (1:200, Abcam, #AB2254) in blocking solution (PBS with 5% BSA) for 1.5 hours at RT in a humid chamber. After washing with PBS, cells were incubated with Alexa Fluor 555–conjugated anti-rabbit antibody (1:1000, Cell Signaling Technology) for 45 min at RT in a humid, light-protected chamber, and then stained with DAPI (1 μg/mL in PBS) for 5 min at RT in a humid, light-protected chamber. Images were acquired using cellSens Imaging Software (Olympus) and merged using ImageJ. AURKB intensity per nucleus was quantified using ImageJ.

### Drug treatment

Cells were seeded in 96-well plates using the Multidrop™ Combi Reagent Dispenser (ThermoFisher). 24 hours later, cells were treated with the drugs of interest. Cell viability was measured after 72 hours of drug treatment using the MTT assay (Sigma, M2128), with 500 μg/mL salt diluted in complete medium and incubated at 37°C for 3 hours. Formazan crystals were solubilized using 10% Triton X-100 and 0.1N HCl in isopropanol, and plates were covered with foil and shaken at 200 rpm for 2 hours. Absorbance was measured at 570 nm and 630 nm. The area under the curve (AUC) for each drug was calculated using GraphPad PRISM 10. Details of all drugs are available in Supplementary Table S1.

### Live-cell imaging using LiveCyte®

Live-cell imaging was performed using LiveCyte® (Phase Focus). Cells were seeded in glass-bottom, imaging-compatible 96-well plates (Corning) at a density of 2,000 cells per well for CAL51, and 8,000 cells per well for MCF7 and MDA-MB cell lines. For proliferation analysis, images were acquired every 2 hours using 10× magnification for 72 hours with phase contrast only. For mitotic aberration analysis, images were acquired every 5 min using 20× magnification for 48 hours with both phase contrast and GFP channels.

### Flow Cytometry analyses

For cell cycle analysis in MCF7 cells, 200,000 cells per well were seeded in 6-well plates and treated for 48 hours with 5nM barasertib or DMSO (control). Cells were then collected and fixed with ice-cold 70% ethanol for 30 min on ice. Ethanol was washed off, and cells were stained with 50 µg/mL propidium iodide (BioLegend) and 0.1 mg/mL RNase A (Invitrogen) in PBS for 15 min at RT. Flow cytometry acquisition was performed on a CytoFLEX® (Beckman Coulter), and data were analyzed using CytExpert v2.4 software (Beckman Coulter). Gating of live cells and singlets was common across all analyzed samples, whereas cell cycle phase gating was determined individually for each cell line.

For cell cycle analysis in CAL51 cells, 50,000 cells per well were seeded in 6-well plates and treated for 48 hours with 10nM barasertib or DMSO (control). Cells were stained with 1 ng/mL Hoechst 33342 (ThermoFisher) for 1 hour, washed with PBS, and collected for analysis. Flow cytometry acquisition was performed on a CytoFLEX® (Beckman Coulter), and data were analyzed using Kaluza v2.1 software (Beckman Coulter). Gating of live cells and singlets was common across all analyzed samples, whereas WGD-positive cells were determined individually per cell line. Examples of the gating strategy are shown in Supplementary Fig. S12.

For cell death analysis in CAL51, 50,000 cells per well were seeded in 6-well plates and treated with 10nM barasertib or DMSO (control) for 48 hours. Cells were stained with Pacific Blue™ Annexin V (BioLegend) according to the manufacturer’s protocol. Flow cytometry acquisition was performed on a CytoFLEX® (Beckman Coulter), and data were analyzed using Kaluza v2.1 software (Beckman Coulter). Gating of live cells and singlets was common across all analyzed samples, whereas Annexin-positive cells were determined individually per cell line.

### Statistical analysis

Statistical analyses were performed using *GraphPad Prism* version 10. For comparisons between two groups, data normality was first assessed using the Shapiro-Wilk test. If the data did not follow a normal distribution, a Mann-Whitney U test was used. When the data followed a normal distribution, equality of variance was evaluated; if variances were equal, an unpaired two-tailed Student’s t-test was used, and if unequal, a Welch’s t-test was used. All tests were two-tailed unless a directional hypothesis had been established *a priori*, in which case a one-tailed test was used. When a single value was normalized to 1, a one-sample t-test was used.

For comparisons among more than two groups, if the data followed a normal distribution, a repeated-measures one-way ANOVA followed by Holm-Sidak’s post hoc test was used. If the data did not follow a normal distribution, a Kruskal-Wallis test followed by Dunn’s post hoc test was used. For categorical variables, Chi-square test was used, followed by Holm-Sidak’s post hoc test for multiple comparisons. Kaplan-Meier analysis was used to generate survival curves for overall survival (OS) and relapse-free survival (RFS), and curves were compared using the log-rank test.

A *P* value<0.05 was considered statistically significant: *P*<0.05 *(*), P<0.01 (**), P< 0.001 (***)*, *P*<0.0001 *(****)*, and *n.s.*, not significant.

## Results

### High Prevalence of Del17p and its Association with Adverse Tumor Characteristics

Analysis of primary tumors from the METABRIC cohort (**1**) revealed that Del17p occurred in ∼44% of breast cancer tumors (**Fig. 1A**), spanning all molecular subtypes, including HER2+, Luminal A/B, Claudin-low, Basal-like and Normal-like (Supplementary Fig. S1A). This widespread prevalence of Del17p across subtypes highlights the relevance of studying the loss of this chromosome-arm in breast cancer (**17**). Tumors harboring Del17p demonstrated significantly worse clinical outcomes, including reduced overall survival (OS) (**Fig. 1B**) and shorter relapse-free survival (RFS) (**Fig. 1C**), and were associated with higher tumor grade (**Fig. 1D**), stage (**Fig. 1E**), and Nottingham Prognostic Index (NPI) (**Fig. 1F**). Del17p was associated with worse overall survival across all molecular subtypes, although this association did not reach statistical significance for the Basal-like and HER2+ subtypes (Supplementary Fig. S1B). These data indicate that Del17p is strongly linked to poor prognosis and more aggressive tumor behavior across the METABRIC cohort.

**Figure 1:**
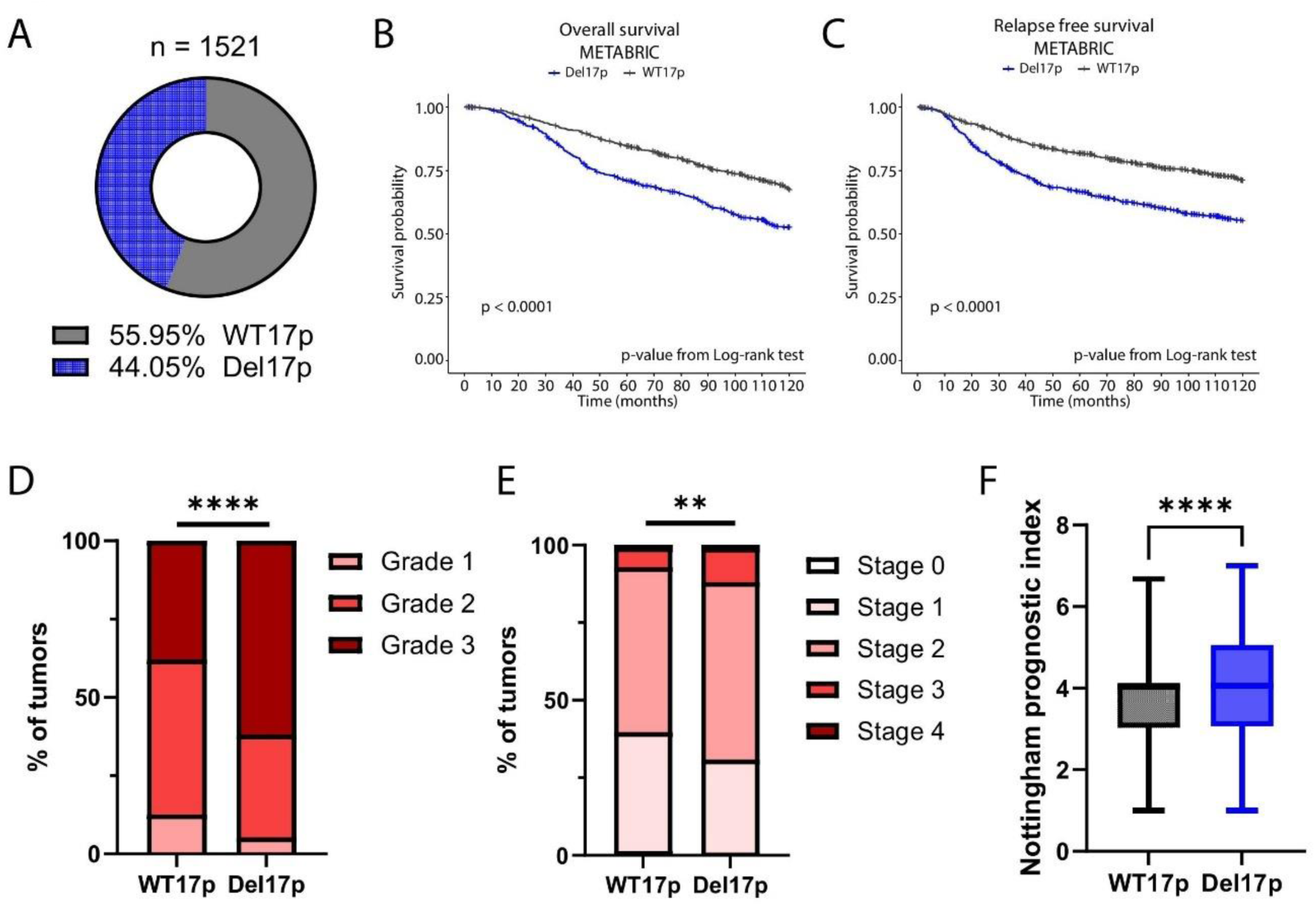
Prevalence of Del17p in breast cancer and its association with tumor features. **A,** Prevalence of Del17p across human primary breast cancer samples from the METABRIC dataset (n=1521). **B,** Comparison of overall survival between patients with Del17p tumors and patients with WT17p tumors, using the METABRIC dataset (n=935). *P*<0.0001 from a Log-rank test. **C,** Comparison of relapse-free survival between patients with Del17p tumors and patients with WT17p tumors, using the METABRIC dataset (n=982). *P*<0.0001 from a Log-rank test. **D,** Comparison of tumor grade between Del17p tumors and WT17p tumors, using the METABRIC dataset (n=1451). *P*<0.0001 from a Chi-squared test. **E,** Comparison of tumor stage between Del17p tumors and WT17p tumors, using the METABRIC dataset (n=1086). *P*=0.0068 from a Chi-squared test. **F,** Comparison of the Nottingham prognostic index (NPI) between Del17p tumors and WT17p tumors, using the METABRIC dataset (n=1449). *P*<0.0001 from a two-tailed Mann-Whitney U-test.

To verify these findings in an independent cohort, we analyzed the TCGA dataset (**24**). Consistent with the METABRIC analysis, Del17p was a common event in the TCGA breast cancer tumors (in ∼56% of patients; Supplementary Fig. S1C), and was associated with shorter disease-free survival (DFS) (Supplementary Fig. S1D), and a higher tumor stage (Supplementary Fig. S1E). Together, these results underscore that Del17p is highly prevalent in breast cancer, where it is associated with aggressive tumor features, making it a desired target for breast cancer therapy.

### Del17p Impacts p53 Activity and is Associated with Tumor Proliferation

We recently performed a genomic analysis of aneuploidy drivers that identified *TP53* as a driver of Del17p across multiple cancer types, including breast cancer (**7**), in line with multiple previous reports (**6, 9, 28-29**). Moreover, in another study we recently identified Del17p-mediated inactivation of *TP53* as a major contributor to breast cancer brain metastasis (**8**). Therefore, we next examined how Del17p affects p53 function, and how it interacts with p53 mutations to affect tumor biology. We stratified the tumors from the METABRIC cohort based on their Chr17p copy number status and p53 mutation and copy-number status (**Fig. 2A**, see Methods). It is known that biallelic inactivation of p53 is often achieved through mutation of one allele and copy number loss of the other (**8, 29**). Indeed, almost a quarter of the METABRIC tumors harbored both p53 mutation and Del17p, indicating biallelic p53 inactivation (**Fig. 2A**). However, in ∼20% of the breast cancer tumors, Del17p was present whereas the other *TP53* allele was intact, and in an additional ∼10% of the tumors, *TP53* was mutant without the chromosome-arm loss (**Fig. 2A**). Tumors representing all four genotypes were present across all molecular subtypes, and Del17p-mediated biallelic inactivation of p53 was particularly common in the basal and HER2+ subtypes (Supplementary Fig. S2A). Gene set enrichment analysis (GSEA) demonstrated that tumors with Del17p exhibited reduced p53 activity in comparison to WT17p tumors, when *TP53* mutation status was not taken into consideration (**Fig. 2B**, left; Supplementary Table S2). Importantly, tumors with Del17p and an intact wild-type *TP53* allele (Del17p;*TP53*-WT) showed significantly lower p53 activity compared to tumors without Del17p (WT17p;*TP53-*WT) (**Fig. 2B**, middle; Supplementary Table S2). As expected, tumors with both Del17p and a *TP53* mutation (Del17p;*TP53*-mut) displayed the strongest reduction in p53 activity (**Fig. 2B**, right; Supplementary Table S2). These findings suggest that the monoallelic disruption of *TP53* due to Del17p is sufficient to impair p53 function and downregulate the p53 pathways, which is further exacerbated by mutations in the remaining allele, in agreement with our recent analysis (**8**).

**Figure 2:**
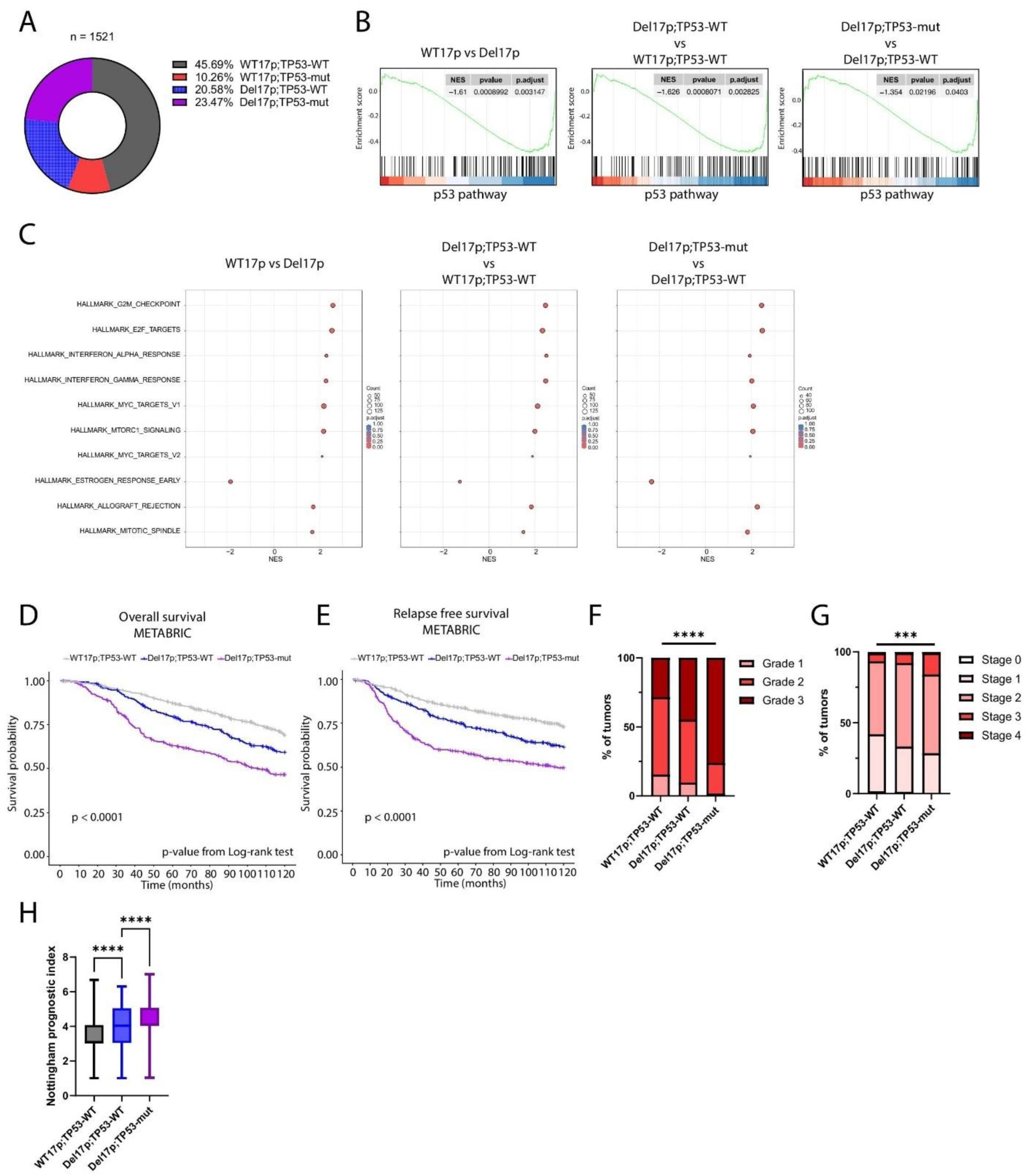
Del17p is associated with p53 pathway downregulation and increased tumor aggressiveness. **A,** Prevalence of Del17p and *TP53* mutation in human primary breast cancer samples from the METABRIC dataset (n=1521) (see Material and Methods). **B,** Gene set enrichment analysis (GSEA) of the METABRIC dataset, showing downregulation of the p53 pathway gene set (P53_DN_V2_DN from the ‘Oncogenic signatures’ dataset, c6) in: tumors with Del17p compared to tumors with WT17p (left); tumors with Del17p;*TP53*-WT compared to tumors with WT17p;*TP53*-WT (middle); tumors with Del17p;*TP53*-mut compared to tumors with Del17p;*TP53*-WT (right). NES=-1.61, *Q*=0.003; NES=-1.62, *Q*=0.002; and NES=-1.35, *Q*=0.04, respectively. **C,** GSEA of the METABRIC dataset, showing enrichment for cell cycle-related pathways (e.g., G2M checkpoint, E2F targets) in: tumors with Del17p compared to tumors with WT17p (left); tumors with Del17p;*TP53*-WT compared to tumors with WT17p;*TP53*-WT (middle); tumors with Del17p;*TP53*-mut compared to tumors with Del17p*TP53*-WT (right). Plots show enrichment for the ‘Hallmark’ gene sets. **D,** Comparison of overall survival between METABRIC patients stratified by their Del17p and *TP53* mutation status, showing worse survival in the patients with Del17p;*TP53*-WT tumors in comparison to those with WT17p;*TP53*-WT tumors (*P*=0.0015), and the worst survival in the patients with Del17p;*TP53*-mut tumors (*P*=0.0002 when compared to the Del17p;*TP53*-WT tumors). n=1255 tumors, P-values from a Log-rank test. **E,** Comparison of relapse-free survival between METABRIC patients stratified by their Del17p and *TP53* mutation status, showing worse survival in the patients with Del17p;*TP53*-WT tumors in comparison to those with WT17p;*TP53*-WT tumors (*P*=0.0002), and the worst survival in the patients with Del17p;*TP53*-mut tumors (*P*=0.0002 when compared to the Del17p;*TP53*-WT tumors). n=1365 tumors, P-values from a Log-rank test. **F,** Comparison of tumor grade between METABRIC patients stratified by their Del17p and *TP53* mutation status, showing a higher tumor grade in the Del17p;*TP53*-WT tumors compared to WT17p;*TP53*-WT and the highest tumor grade in the Del17p;*TP53*-mut tumors. n=1297 tumors, *P*<0.0001 from a Chi-squared test. **G,** Comparison of tumor stage between METABRIC patients stratified by their Del17p and *TP53* mutation status, showing a higher tumor stage in the Del17p;*TP53*-WT tumors compared to WT17p;*TP53*-WT and the highest tumor stage in the Del17p;*TP53*-mut tumors. n=978 tumors, *P*=0.0003 from a Chi-squared test. **H,** Comparison of the Nottingham Prognostic Index (NPI) score between METABRIC patients stratified by their Del17p and *TP53* mutation status, showing a worst prognostic index in the Del17p;*TP53*-WT tumors compared to WT17p;*TP53*-WT (*P*<0.0001), and the worst prognostic index in the Del17p;*TP53*-mut tumors (*P*<0.0001 when compared to Del17p;*TP53*-WT). n=1300 tumors, P-value from a one-way Kruskal-Wallis test with Dunn’s correction for multiple comparisons.

Consistent with the reduction in p53 activity, GSEA revealed that Del17p tumors were characterized by upregulation of cell cycle-related pathways, such as the G2M checkpoint, MYC targets and E2F targets (**Fig. 2C**, left panel; Supplementary Fig. S3A, left panel; Supplementary Table S2). These pathways, which are key drivers of tumor proliferation, were also upregulated when Del17p;*TP53*-WT tumors were compared to WT17p;*TP53*-WT tumors (**Fig. 2C**, middle panel; Supplementary Fig. S3A, middle panel; Supplementary Table S2), and they were most strongly upregulated in Del17p;*TP53*-mut tumors (**Fig. 2C**, right panel; Supplementary Fig. S3A, right panel; Supplementary Table S2). The same pattern was observed in the TCGA cohort, where Del17p tumors elevated multiple transcriptional programs of cell proliferation (Supplementary Fig. S3B and S3C; Supplementary Table S2).

Further analysis of tumor clinical features from the METABRIC cohort revealed that Del17p;*TP53*-WT tumors had a worse prognosis and were associated with more advanced tumor features in comparison WT17p;*TP53*-WT tumors, while the poorest prognosis and most advanced tumor features were observed in Del17p;*TP53*-mut tumors (**Fig. 2D-H**; Supplementary Fig. S2B). The same trend was observed when analyzing the breast cancer tumors from the TCGA cohort (Supplementary Fig. S2C-S2E). Together, these results indicate that Del17p impairs p53 pathway activity and promotes uncontrolled proliferation, thereby contributing to tumor aggressiveness and to cancer progression.

### AURKB is a Selective Vulnerability of Del17p Breast Cancer Cells

Previous studies have identified genes that become more essential for cancer cells following copy number loss of one of their alleles, giving rise to hemizygous dependencies. Cancer cells have shown to be more dependent on such CYCLOPS genes, highlighting them as therapeutic targets (**9, 14-15**). Notably, Del17p was previously shown to sensitize colorectal and breast cancer cells to targeting of the Chr17p-residing RNA polymerase, POLR2A (**17, 30**), as well as its activator, RBX1, in prostate cancer (**16**). Here, we set out to identify novel hemizygous dependencies in Del17p breast cancer cells. As expected, Del17p tumors exhibited a significant reduction in the expression levels of multiple genes located on Chr17p, both in the METABRIC and in the TCGA datasets (Supplementary Fig. S4A and S4B), suggesting a broad effect of Del17p on the expression of Chr17p-residing genes. These results highlight the potential to target additional Chr17p genes that play a role in tumor fitness and are vulnerable due to hemizygosity.

To identify such genes, we performed an unbiased analysis of gene expression and functional dependencies across 65 human breast cancer cell lines, using the DepMap dataset (**27**) (**Fig. 3A**). Specifically, we searched for genes whose mRNA and protein expression levels are reduced upon Del17p, and that Del17p cell lines are more sensitive to their genetic depletion by RNAi and/or by CRISPR-Cas9 (**Fig. 3B**; Supplementary Table S3; see Materials and Methods). Among these genes, of particular interest are those that are <u>not</u> downregulated and are <u>not</u> more essential in cells that have lost *TP53* <u>without</u> Del17p (**Fig. 3B**; Supplementary Table S3; see Materials and Methods). The top ‘hit’ of this analysis was the important mitotic kinase, AURKB (**Fig. 3B**). Del17p cell lines exhibited reduced AURKB mRNA and protein expression levels, independent of their *TP53* mutation status (**Fig. 3C**; Supplementary Fig. S5A and S5B). These findings held true when we controlled for several potential confounders: the *TP53* mutation status, the copy number status of chromosome-arm 17q, the aneuploidy score, the doubling time, and the breast cancer molecular subtypes (Supplementary Fig. S5A and S5B; see Materials and Methods). Importantly, in both METABRIC and TCGA datasets, Del17p tumors consistently showed lower AURKB expression levels, independent of *TP53* mutation status (**Fig. 3D**), demonstrating that Del17p results in AURKB downregulation in human tumors as well. Furthermore, AURKB was the most differentially essential Chr17p-residing gene in both *TP53*-WT and *TP53*-mutant cell lines: in both cases, Del17p breast cancer cell lines were more sensitive to the genetic depletion of AURKB in comparison to WT17p cell lines (**Fig. 3E**). These findings also held true when controlling for all of the abovementioned potential confounders (Supplementary Fig. S6; Supplementary Fig. S7; see Materials and Methods), substantiating AURKB as a Del17p-associated dependency of human breast cancer cells.

**Figure 3:**
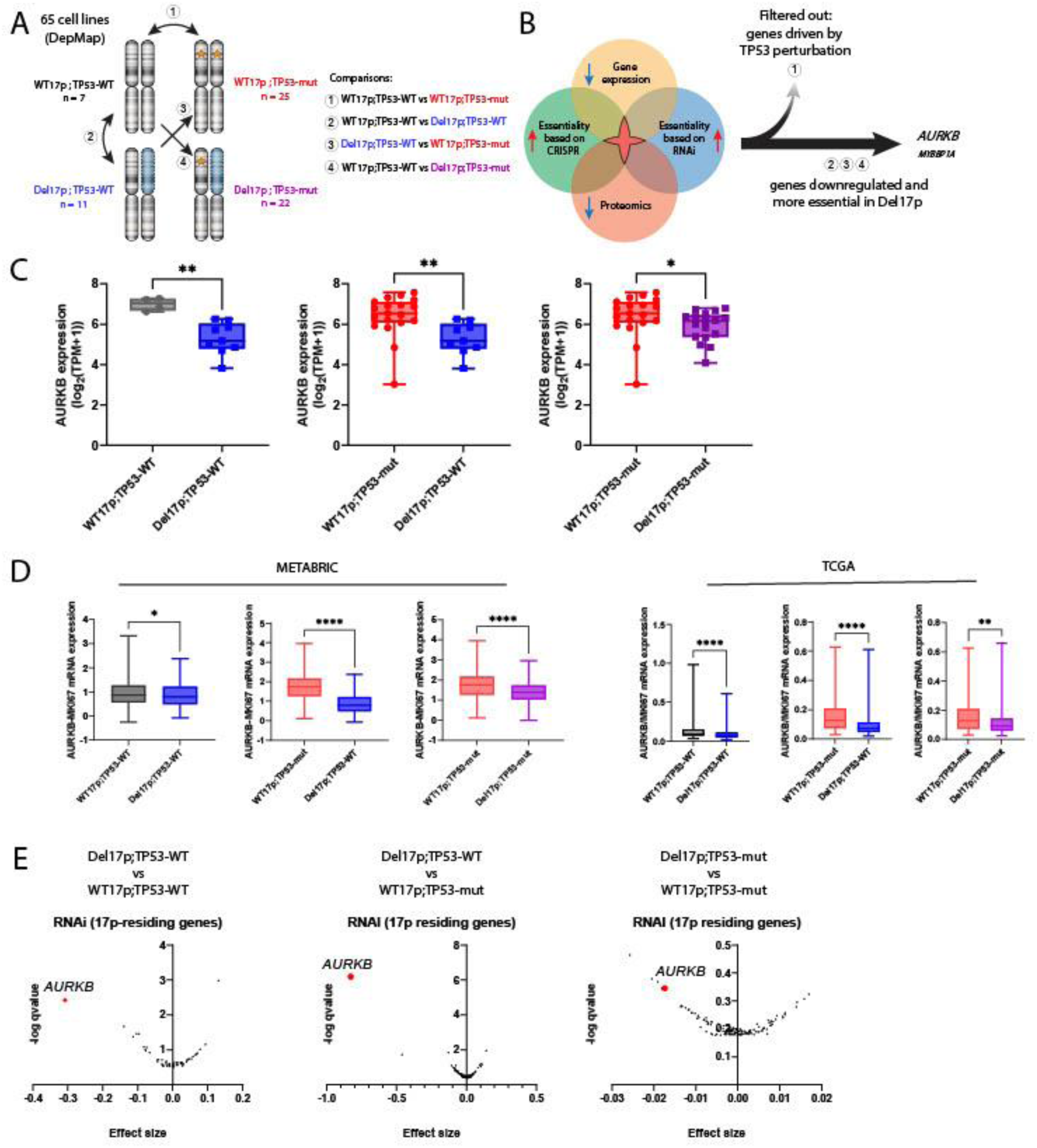
Del17p is associated with reduced expression of, and increased dependency on, AURKB. **A,** Schematic illustration of the stratification of 65 human breast cancer cell lines in the DepMap dataset by Chr17p status and *TP53* mutation status. Arrows and numbers indicate paired groups that were used for the downstream analyses. **B,** Venn diagram illustrating the intersection of analyses across various datasets, highlighting genes that are both downregulated and more essential in Del17p compared to WT17p breast cancer cell lines, but are not downregulated or more essential in *TP53*-mut vs. *TP53*-WT cell lines. **C,** Comparison of AURKB mRNA levels in breast cancer cell lines shows that Del17p is associated with its reduced expression. WT17p;*TP53*-WT compared to cell lines with Del17p;*TP53*-WT, *P*=0.0019 from a two-tailed unpaired Student’s t-test. WT17p;*TP53*-mut compared to cell lines with Del17p;*TP53*-WT, *P*=0.0029 and WT17p;*TP53*-mut compared to cell lines with Del17p;*TP53*-mut, *P*=0.0252. P-values from a two-tailed Mann-Whitney U-test. **D,** Comparisons of AURKB mRNA levels in breast cancer clinical cohorts, METABRIC (left) and TCGA (right), show that Del17p is associated with reduced gene expression of AURKB. METABRIC data: WT17p;*TP53*-WT compared to cell lines with Del17p;*TP53*-WT, *P*=0.0287. WT17p;*TP53*-mut compared to cell lines with Del17p;*TP53*-WT, *P*<0.0001. P-values from a two-tailed Mann-Whitney U-test. WT17p;*TP53*-mut compared to cell lines with Del17p;*TP53*-mut, *P*<0.0001 from a two-tailed unpaired Welch’s t-test. TCGA data: *P*<0.0001, *P*<0.0001, *P*=0.0012, respectively. P-values from a two-tailed Mann-Whitney U-test. AURKB mRNA levels were normalized to MKI67 mRNA levels to control for proliferation rate (as it is strongly correlated with AURKB expression levels). **E,** Comparison of the sensitivity of human breast cancer cell lines to the knockdown of Chr17p-residing genes, based on the Novartis RNAi screen (**42**), showing AURKB as the most preferentially essential gene in Del17p cell lines. Shown is the comparison between Del17p;*TP53*-WT vs. WT17p;*TP53*-WT (left); Del17p;*TP53*-WT vs. WT17p;*TP53*-mut (middle); Del17p;*TP53*-mut vs. WT17p;*TP53*-mut (right).

To experimentally validate that Del17p is associated with increased vulnerability to AURKBi, we compared two p53-deficient basal-like breast cancer cell lines, MDA-MB-231 and MDA-MB-468, which differ in their Chr17p copy number status (**27**). MDA-MB-468 cells, which harbor Del17p, exhibited significantly lower AURKB expression (**Fig. 4A-C**; Supplementary Fig. S8A and S8B) and increased sensitivity to barasertib (**Fig. 4D** and **E**), a potent and selective AURKB inhibitor (**31**), compared to the WT17p MDA-MB-231 cells. To rule out that the difference in drug sensitivity was due to an increased resistance of MDA-MB-231 cells to drugs in general, we compared the response of the two cell lines to four other anti-cancer drugs, and found that barasertib exhibited the largest and most significant differential sensitivity of all (Supplementary Fig. S8C and S8D). Live-cell imaging confirmed that barasertib treatment impaired the proliferation of MDA-MB-468 cells more than that of MDA-MB-231 cells (**Fig. 4F** and **G**). These results suggest that Del17p tumor cells might be particularly sensitive to AURKBi.

**Figure 4:**
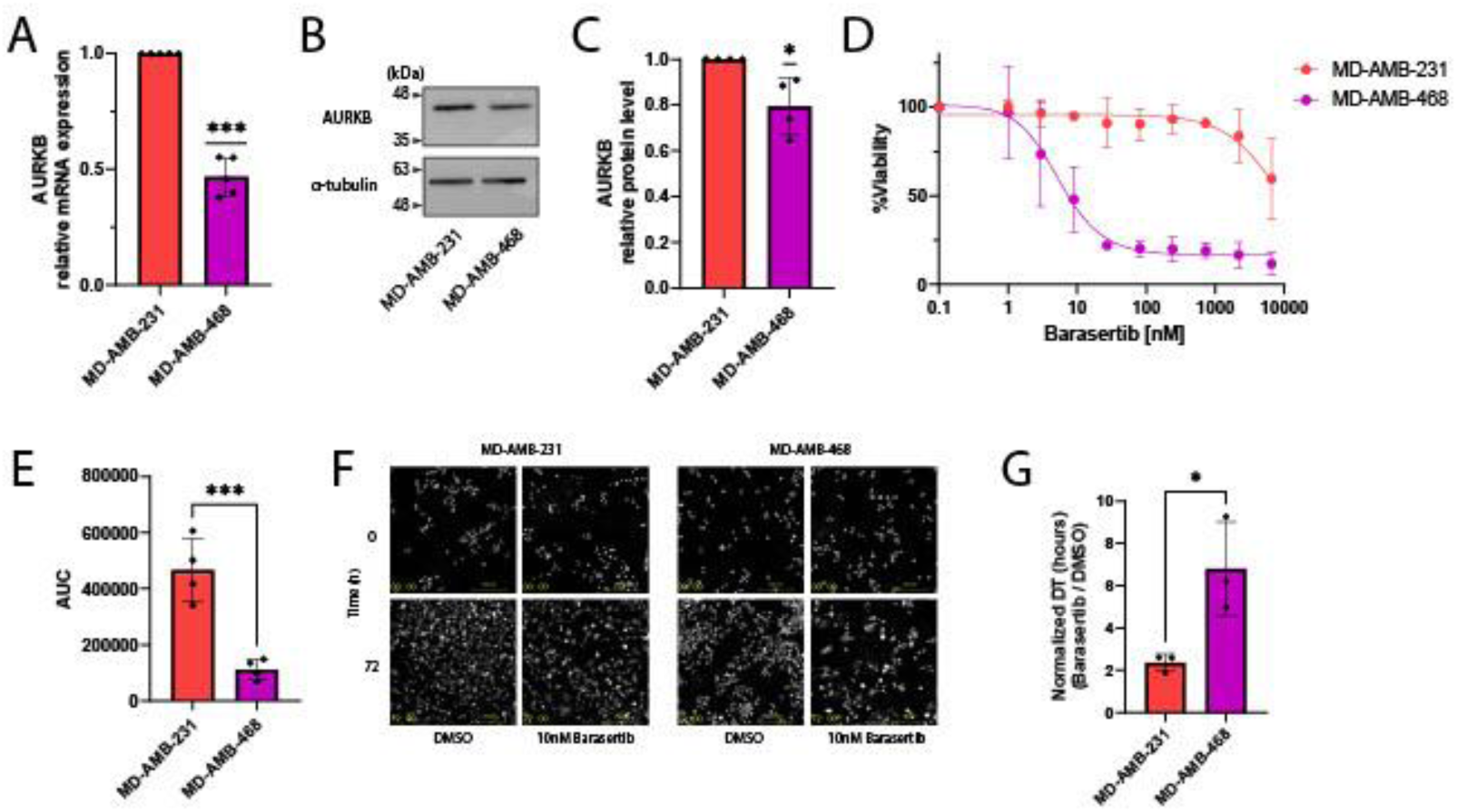
Del17p is associated with reduced AURKB expression and higher sensitivity to its inhibition in human breast cancer cell lines. **A,** Comparison of AURKB mRNA levels between MD-AMB-231 and MD-AMB-468 cells, using RT-qPCR. *P*=0.0002 from a one-sample t-test. n=5 biological repeats. **B,** Comparison of AURKB protein levels between MD-AMB-231 and MD-AMB-468 cells, using WB. α-tubulin used as a control. **C,** Quantification of the WB results shown in (C). *P*=0.0466 from a one-sample t-test. n=4 biological repeats. **D,** Dose-response curves comparing the response of MD-AMB-231 and MD-AMB-468 cells to barasertib. Cells were exposed to the drug for 72 hours. Each curve shows the average of 3 biological repeats. **E,** Comparison of the Area Under the Curve (AUC) values from the experiments shown in (D). *P*=0.0005 from a one-tailed unpaired Student’s t-test. n=3 biological repeats. **F,** Representative images of MD-AMB-231 and MD-AMB-468 cells exposed to 10nM barasertib, before and after 72 hours of drug exposure. **G,** Live-cell imaging-based comparison of cell proliferation between MD-AMB-231 and MD-AMB-468 cells exposed to 10nM barasertib for 72 hours. *P*=0.05 from a one-tailed Mann-Whitney U-test. n=3 biological repeats. DT=doubling time.

### Validation and Characterization of AURKB Sensitivity in MCF7 Cells with Del17p

We extended these observations by investigating AURKB expression and dependency in a semi-isogenic system based on MCF7, a p53-competent human luminal breast cancer cell line (**32**). Importantly for the current study, different MCF7 cultures differed in their Chr17p copy number state. We selected 3 WT17p strains (strains D,E,K) and 3 Del17p strains (strains L,O,N) for comparisons (**Fig. 5A**). Consistent with the findings in the p53-deficient basal MDA-MB-231 and MDA-MB-468 cell lines, Del17p MCF7 strains showed reduced AURKB mRNA and protein expression levels in comparison to WT17p MCF7 strains (**Fig. 5B-D**; Supplementary Fig. S9A and S9B).

**Figure 5:**
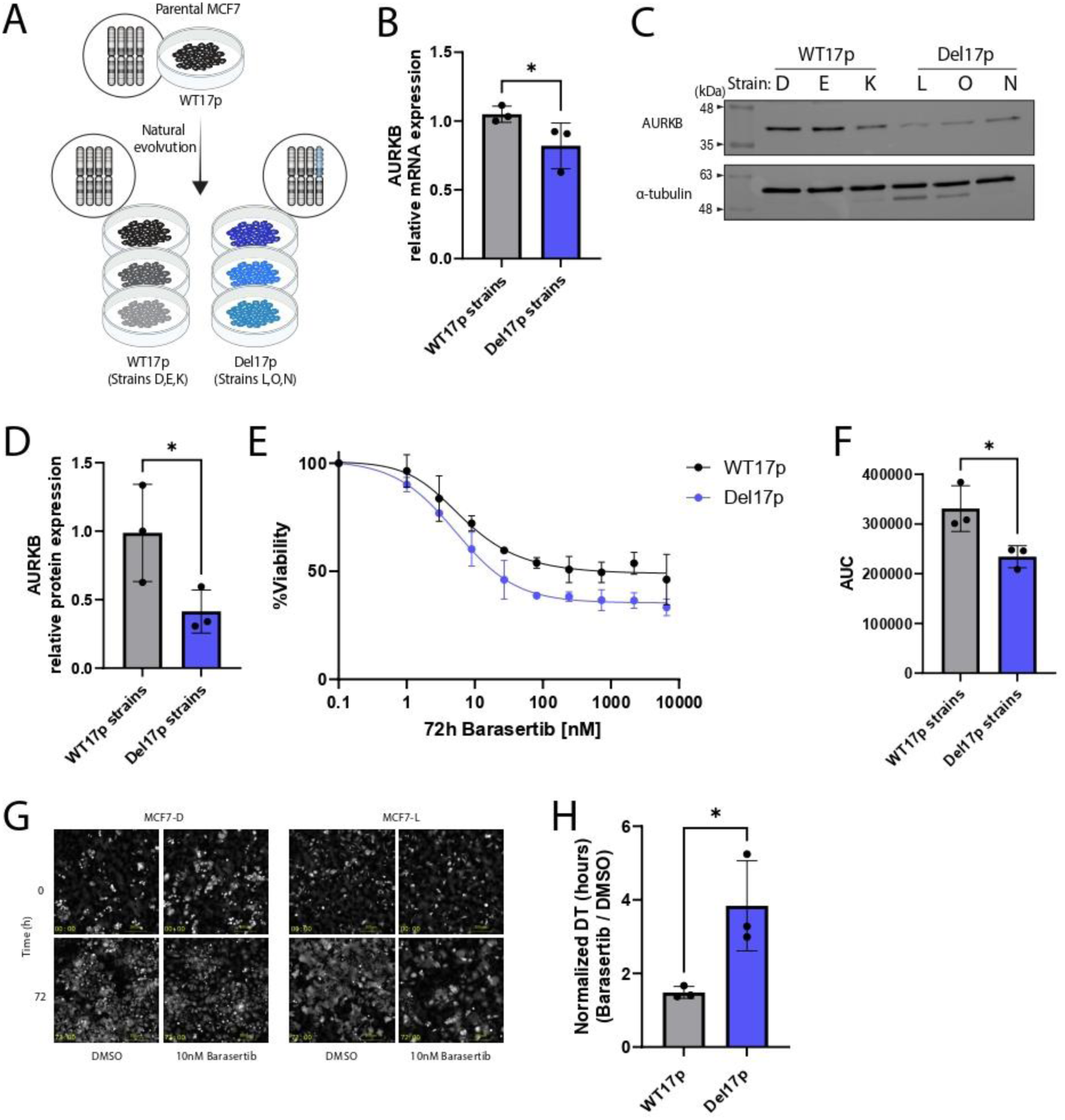
Del17p reduces AURKB expression and increases sensitivity to barasertib in MCF7 cells. **A,** Schematic illustration of the in-vitro evolution of the MCF7 strains. The semi-isogenic MCF7 system, comprising three WT17p and three Del17p MCF7 strains along with their copy number status. **B,** Comparison of AURKB mRNA levels between WT17p and Del17p MCF7 strains, using RT-qPCR. The mean expression of each strain was calculated from 5 biological repeats. *P*=0.0437 from a one-tailed unpaired Student’s t-test; n=3 strains per group. **C,** Comparison of AURKB protein levels between WT17p and Del17p MCF7 strains, using WB. α-tubulin used as a control. **D,** Quantification of the WB results shown in (C). The mean expression of each strain was calculated from 4 biological repeats. *P*=0.0484 from a one-tailed unpaired Student’s t-test. n=3 strains per group. **E,** Dose-response curves comparing the response of WT17p and Del17p MCF7 cells to barasertib. Each curve shows the average of 3 biological repeats. **F,** Comparison of the Area Under the Curve (AUC) values from the experiments shown in (E). *P*=0.0151 from a one-tailed unpaired Student’s t-test. n=3 biological repeats. **G,** Representative live-cell imaging photos of a WT17p strain (MCF7-D) and a Del17p strain (MCF7-L) treated with 10nM barasertib for 72 hours. **H,** Live-cell imaging-based comparison of cell proliferation between WT17p MCF7 strains and Del17p MCF7 strains exposed to 10nM barasertib for 72 hours. *P*=0.0385 from a one-tailed unpaired Welch’s t-test. n=3 biological repeats.

Next, we compared the drug response between WT17p and Del17p MCF7 strains. First, we recapitulated in our system the known Del17p-induced cellular dependency on POLR2A. Indeed, the Del17p MCF7 strains were more sensitive to the drug α-amanitin, which inhibits POLR2A (Supplementary Fig. S9C and S9D), in line with previous studies in other isogenic systems (**16-17**), demonstrating the suitability of our system for identifying Del17p-associated cellular vulnerabilities. Next, we re-analyzed drug screen data performed on our 6 strains (**32**) and found that the drug that was most differentially-effective in the Del17p strains was the Survivin inhibitor YM-155 (Supplementary Fig. S9E and S9F). We then recapitulated this result experimentally (Supplementary Fig. S9G and S9H). This finding is of great interest since Survivin directly interacts with AURKB as part of the CPC (**18**), suggesting that the increased sensitivity of Del17p cells to this drug may also result from AURKB hemizygosity. Lastly, we treated the MCF7 strains with barasertib and found that the Del17p strains were significantly more sensitive to AURKBi than their WT17p counterparts (**Fig. 5E** and **F**; Supplementary Fig. S9I), in line with our findings in the MDA-MB cell lines (**Fig. 4**). Live-cell imaging further confirmed that barasertib treatment reduced cell proliferation to a greater extent in the Del17p MCF7 strains (**Fig. 5G** and **H**).

To further explore the molecular mechanism underlying the increased sensitivity of Del17p breast cancer cells to AURKBi, we examined the consequences of barasertib on mitotic progression, with a focus on whole-genome doubling (WGD), a well-known consequence of AURKBi (**33**). Flow cytometry analysis revealed that barasertib-treated Del17p MCF7 cells were significantly more likely to undergo WGD compared to WT17p MCF7 cells (**Fig. 6A** and **B**). Live-cell imaging corroborated these findings, showing a significant increase in mitotic aberrations in Del17p cells upon treatment with barasertib (**Fig. 6C** and **D**). Tracking cell fate under barasertib treatment revealed that the majority of mitotic aberrations were mitotic failures, defined as unsuccessful mitoses resulting in WGD (**Fig. 6D-F**): mitotic slippage was characterized by chromosome condensation followed by decondensation without cytokinesis, leading to a mononuclear cell, whereas cytokinesis failure led to incomplete cell division and the formation of a binucleated cell – both resulting in WGD, consistent with our flow cytometry data (**34-35**) (**Fig. 6F**).

**Figure 6:**
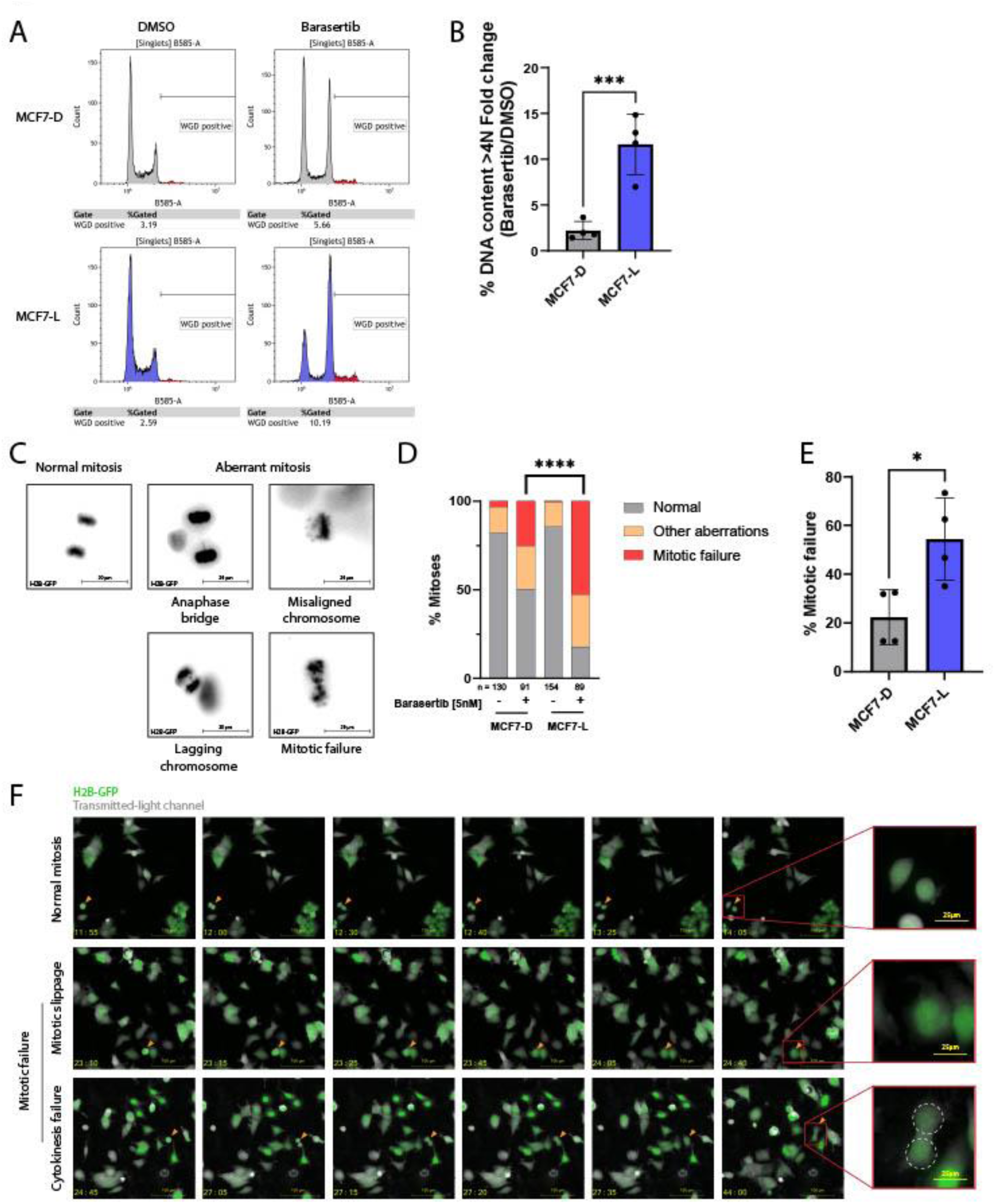
Del17p increases whole-genome doubling in response to AURKB inhibition. **A,** Representative images from a flow cytometry-based cell cycle analysis of WT17p MCF7 cells (MCF7-D) vs. Del17p MCF7 cells (MCF7-L) exposed to either DMSO (control) or 5nM barasertib for 48 hours. **B,** Quantification of the % WGD-positive cells with DNA content greater than 4N. *P*=0.0008 from a one-tailed unpaired Student’s t-test. n=4 biological repeats. **C,** Representative live-cell images of MCF7-L cells stably expressing histone H2B-GFP (black) undergoing normal and aberrant mitoses. **D,** Time-lapse microscopy-based quantification of various types of mitotic aberrations in WT17p MCF7 cells (MCF7-D) and Del17p MCF7 cells (MCF7-L) following exposure to 5nM barasertib. Quantified aberration types were anaphase bridges, misaligned chromosomes, lagging chromosomes and mitotic failures. Statistical comparison was performed between the frequency of normal mitoses and the total frequency of aberrant mitoses (all types combined). *P*<0.0001, from a Chi-squared test. Data represents pooled results, n=4 biological repeats. **E,** Comparison of the fraction of mitotic failure events between WT17p MCF7 cells (MCF7-D) and Del17p MCF7 cells (MCF7-L) exposed to 5nM barasertib for 48 hours. Each point represents one biological repeat, calculated as the number of mitotic failures divided by the total mitoses within that replicate. *P*=0.0143 from a one-tailed unpaired Mann-Whitney U-test. n=4 biological repeats. **F,** Representative time-lapse microscopy images of an MCF7-L cell stably expressing histone H2B-GFP (green). The orange arrow indicates cells undergoing: normal mitosis (top); mitotic slippage, defined as chromosome condensation followed by decondensation without cytokinesis, resulting in a mononucleated cell (middle); and cytokinesis failure, leading to the formation of a binucleated cell indicated by white dashed circles (bottom). Cells were treated with 5nM barasertib. Time (yellow) indicates hours from the start of drug exposure.

Together, these data demonstrate that the Del17p-mediated reduction of AURKB expression creates a higher sensitivity of Del17p cells to further inhibition of AURKB mitotic function, resulting in greater drug-induced mitotic aberrations and cytokinesis failure in Del17p cells. Importantly, this differential dependency holds true in both *TP53*-WT and *TP53*-mutant breast cancer cells, and across breast cancer molecular subtypes.

### AURKB Hemizygosity Confers Sensitivity to AURKBi in a p53-dependent Manner

To explore whether it is indeed the hemizygosity of AURKB that underlies the increased sensitivity of Del17p breast cancer cells to AURKBi, we generated heterozygous knockouts (KO) of *AURKB* (*AURKB*^+/−^) in *TP53*-WT, diploid CAL51 breast cancer cells and validated the heterozygous KO via Sanger sequencing (**Fig. 7A-C**). Single-cell cloning of CAL51 cells following CRISPR-Cas9 targeting of *AURKB* gave rise to 42 single-cell-derived clones. In line with *AURKB* being a ‘common essential gene’ (**27**), we were able to obtain heterozygous *AURKB*-KO – but no homozygous *AURKB*-KO – clones (Supplementary Fig. S10A). RT-qPCR, immunofluorescence and Western blot (WB) analyses confirmed a significant reduction in AURKB mRNA and protein expression levels in three selected heterozygous *AURKB*-KO clones, compared to clones in which *lacZ* was targeted as a control (**Fig. 7D-F**; Supplementary Fig. S10B-S10F). We then subjected the clones to Sanger sequencing, which confirmed that all three harbor exon 2 exclusion in one allele whereas the other allele is intact (Supplementary Fig. S10G).

**Figure 7:**
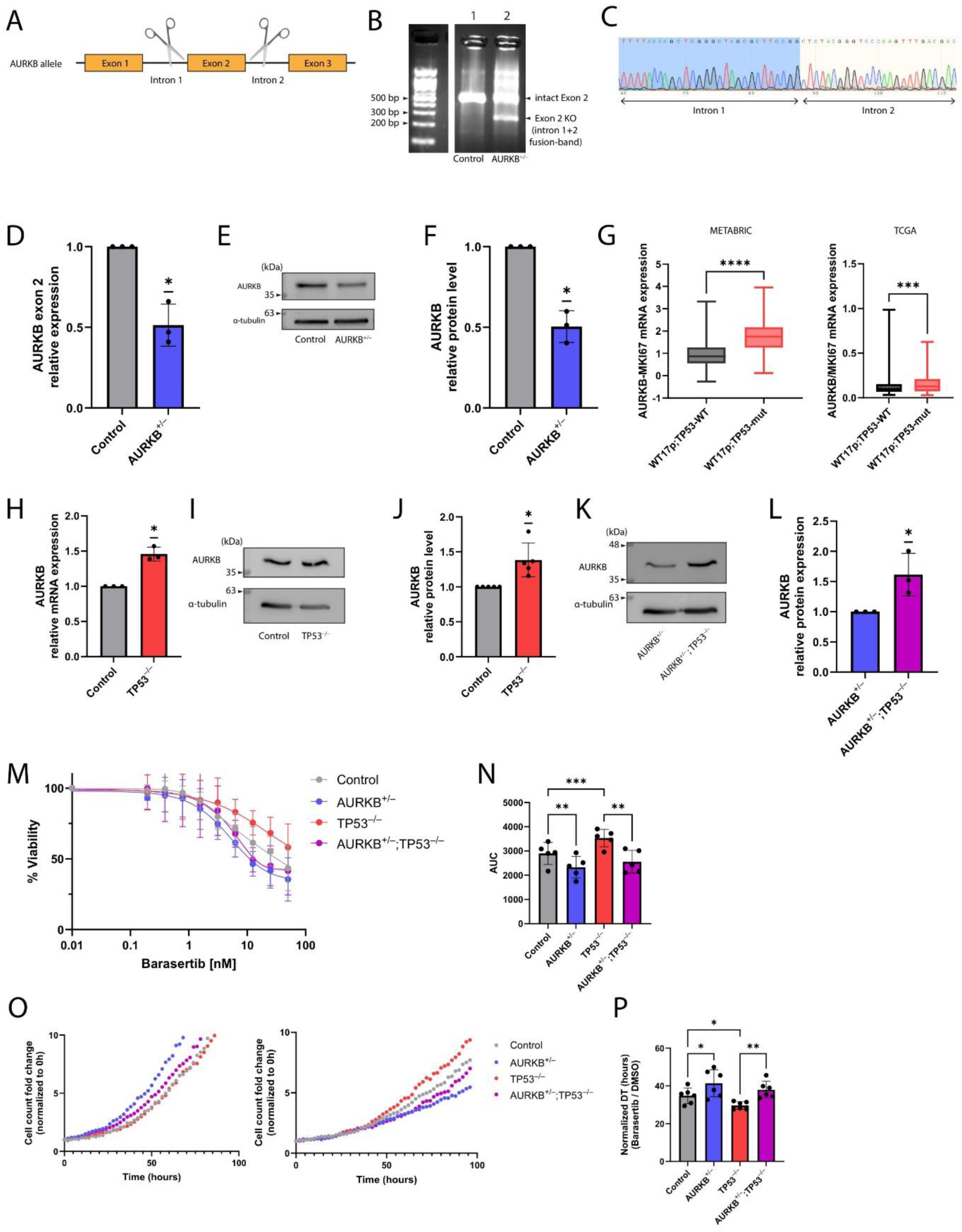
*AURKB* hemizygosity confers sensitivity to AURKB inhibition in a p53-dependent manner. **A,** Schematic Illustration of the strategy applied to generate heterozygous *AURKB*-KO clones, where flanking introns of exon 2, which contains the translation initiation site, were targeted. **B,** Bulk-level PCR-based amplification of *AURKB* exon 2 in CAL51 cells was performed using primers targeting introns 1 and 2. Lane 1 represents cells treated with a single-guide RNA targeting *lacZ* (Control), while lane 2 represents cells treated with single-guide RNAs targeting introns 1 and 2, confirming successful bulk-level heterozygous *AURKB*-KO (*AURKB*^+/−^) in the CAL51 cells. **C,** Sanger sequencing of the fusion PCR product from a bulk CAL51 cell population. Primers flanking the expected fusion junction between introns 1 and 2 were used, demonstrating the fusion of the sequence proximal to the first cut site with the sequence distal to the second cut site. **D,** Comparison of *AURKB* mRNA levels between Control and heterozygous *AURKB*-KO CAL51 cells (a mix of 3 single-cell-derived clones) using RT-qPCR. *P*=0.0231 from a one-sample t-test. n=3 biological repeats. **E,** Comparison of AURKB protein levels between Control and heterozygous *AURKB*-KO CAL51, using WB. α-tubulin used as a control. **F,** Quantification of the WB results shown in (E). *P*=0.0128 from a one-sample t-test. n=3 biological repeats. **G,** Comparison of *AURKB* mRNA expression levels between WT17p;*TP53*-WT and WT17p;*TP53*-mut primary breast cancer samples in METABRIC (left, *P*<0.0001) and TCGA (right, *P*=0.001). P-values from a one-tailed Mann-Whitney U-test. **H,** Comparison of *AURKB* mRNA levels between Control and *TP53*-null (*TP53*^−/−^) CAL51 cells, using RT-qPCR. *P*=0.0149 from a one-sample t-test. n=3 biological repeats. **I,** Comparison of AURKB protein levels between control and *TP53*-null CAL51 cells, using WB. α-tubulin used as a control. **J,** Quantification of the WB results shown in (I). *P*=0.0237 from a one-sample t-test. n=5 biological repeats. **K,** Comparison of AURKB protein levels between *TP53*-WT and *TP53*-null heterozygous *AURKB*-KO CAL51 cells (*AURKB^+/−^* and *AURKB*^+/−^;*TP53*^−/−^, respectively), using WB. α-tubulin used as a control. **L,** Quantification of the WB results shown in (K). *P*=0.0157 from a one-sample t-test. n=3 biological repeats. **M,** Dose-response curves comparing the response to barasertib of Control vs. heterozygous *AURKB*-KO CAL51 cells, on a *TP53*-WT or *TP53*-null background. Cells were exposed to the drug for 72 hours. *TP53*-null cells are more resistant to the drug, but *AURKB* hemizygosity increases the sensitivity on both *TP53* backgrounds. Data represents pooled results, n=5 biological repeats. **N,** Comparison of the Area Under the Curve (AUC) values from the experiments shown in (K). *P*=0.0043 (Control vs. *AURKB*^+/−^); *P*=0.0009 (Control vs. *TP53^−^*^/−^); *P*=0.006 (*TP53^−^*^/−^ vs. *AURKB*^+/−^;*TP53^−^*^/−^). P-values from repeated-measures one-way ANOVA with Holm-Sidak’s correction for multiple comparisons; n=5 biological repeats. **O,** Live-cell imaging-based proliferation curves of the four CAL51 isogenic models exposed to DMSO (left) or to 1nM barasertib (right). **P,** Quantification of the doubling time from the experiments shown in (M). *AURKB* hemizygousity increases sensitivity to barasertib, as reflected by reduced proliferation. In contrast, *TP53*-null cells are more resistant to the drug when AURKB is intact. *P*=0.039 (Control vs. *AURKB*^+/−^); *P*=0.0152 (Control vs. *TP53^−^*^/−^); *P*=0.0046 (*TP53^−^*^/−^ vs. *AURKB*^+/−^;*TP53^−^*^/−^). P-values from repeated-measures one-way ANOVA with Holm-Sidak’s correction for multiple comparisons; n=6 biological repeats. DT=doubling time.

As shown in Figure 1, many Del17p tumors also harbor *TP53* mutations in the remaining Chr17p allele. Previous studies have demonstrated that in lung and colon cancers, knocking out *TP53* resulted in increased expression of AURKB (**36-37**), suggesting that *TP53* loss-of-function (LOF) may lead to the upregulation of AURKB. Indeed, in both the METABRIC and TCGA datasets, we found that *TP53*-mutant breast tumors exhibited elevated levels of AURKB (**Fig. 7G**). Therefore, and given the high prevalence of complete *TP53* LOF in Del17p tumors (**38**), we knocked out *TP53* in our control and heterozygous *AURKB*-KO cells in order to study the interaction between *TP53* mutation, AURKB hemizygosity and barasertib sensitivity. *TP53* KO was validated at the population level by RT-qPCR and WB of p53 and of its direct transcriptional target p21 (Supplementary Fig. S10H and S10I). In line with the previous reports and with the breast cancer clinical data, we found that *TP53* KO resulted in a significant increase in mRNA and protein expression levels of AURKB in the CAL51 cells (**Fig. 7H-L**).

These results suggest that the effect of p53 inactivation on AURKB may depend on the mode of p53 inactivation: when *TP53* is mutated or focally deleted, its LOF increases AURKB expression and activity; however, when *TP53* is lost through Del17p, it is invariably associated with AURKB hemizygosity, which decreases AURKB expression and activity, opposing the p53-induced increase. Interestingly, previous studies that focused on point mutations suggested that p53 LOF rendered the cells more resistant to barasertib (**22-23**), although another study reported conflicting results (**36**). To clarify this discrepancy, we evaluated barasertib sensitivity in our isogenic breast cancer cells with/without *AURKB* hemizygosity and with/without *TP53*-KO. Our *TP53*-null cells were indeed more resistant to barasertib than their *TP53*-WT controls (**Fig. 7M-P**). Importantly, however, in both *TP53*-WT and *TP53*-null cells, heterozygous loss of *AURKB* increased the cellular sensitivity to barasertib, as evaluated by a dose-response viability assay (**Fig. 7M** and **N**) and by live-cell imaging (**Fig. 7O** and **P**). In other words, *AURKB* hemizygosity re-sensitized *TP53*-null cells to the drug. Similarly, *AURKB* hemizygosity also rendered the cells sensitive to the Survivin inhibitor YM-155 (Supplementary Fig. S11A and S11B), in line with our results in the MCF7 cells (Supplementary Fig. S9G and S9H), further linking the effect of AURKB hemizygosity on the sensitivity to AURKB inhibition to its important mitotic role as part of the CPC.

Finally, we examined how the interplay between *TP53* LOF and *AURKB* hemizygosity determines the mitotic consequences of AURKBi. Live-cell imaging demonstrated an increased prevalence of mitotic aberrations in the heterozygous *AURKB*-KO cells following treatment with barasertib (**Fig. 8A**). Notably, most of these mitotic aberrations were mitotic failure events (i.e., mitotic slippage or cytokinesis failure) (**Fig. 8B**). Interestingly, *TP53*-null cells with intact *AURKB* exhibited fewer mitotic failure events than *TP53*-WT cells upon barasertib treatment, but this protective effect of *TP53* KO was completely abolished in the context of *AURKB* hemizygosity (**Fig. 8A** and **B**). Congruently, flow cytometry analysis revealed increased prevalence of barasertib-induced WGD (**Fig. 8C** and **D**; Supplementary Fig. S12) and apoptosis (**Fig. 8E**; Supplementary Fig. S12) in the hemizygous *AURKB* cells. Importantly, *TP53*-null cells with intact *AURKB* were more resistant to the drug, displaying few WGD events and apoptotic cells, whereas heterozygous *AURKB*-KO enhanced drug-induced WGD and apoptosis in both *TP53*-WT and *TP53*-null backgrounds (**Fig. 8C-E**).

**Figure 8:**
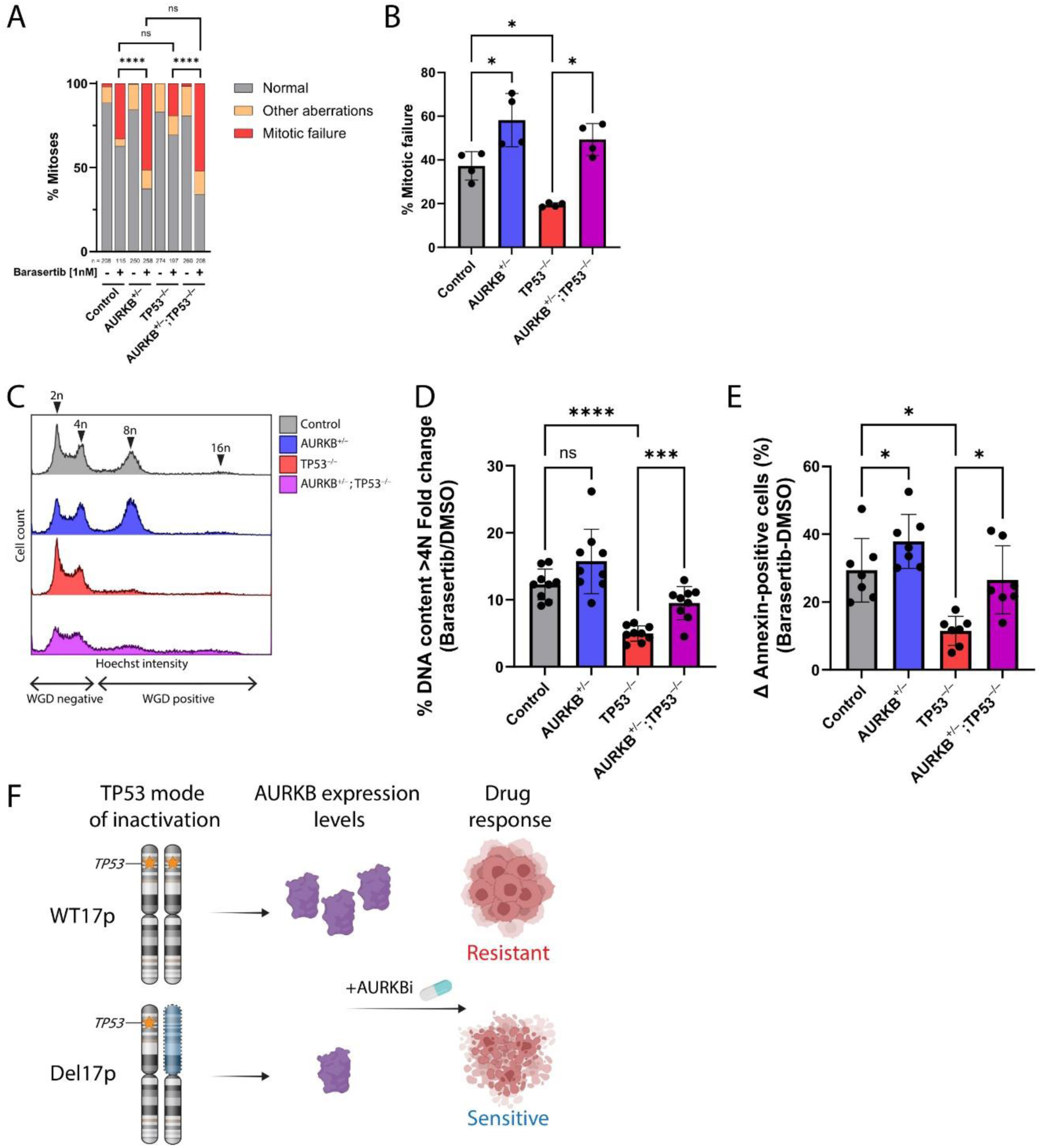
*AURKB* hemizygosity increases whole-genome doubling and apoptosis in response to barasertib. **A,** Time-lapse microscopy-based quantification of various types of mitotic aberrations in CAL51 isogenic models following exposure to 1nM barasertib. Quantified aberration types were anaphase bridges, misaligned chromosomes, lagging chromosomes and mitotic failures. Statistical comparison was performed between the frequency of normal mitoses and the total frequency of aberrant mitoses (all types combined). *P*<0.0001 (Control vs. *AURKB*^+/−^ cells, treated with barasertib); *P*=n.s. (Control vs. *TP53^−^*^/−^ cells, treated with barasertib); *P*<0.0001 (*TP53^−^*^/−^ vs. *AURKB*^+/−^;*TP53^−^*^/−^ cells, treated with barasertib); *P*=n.s. (*AURKB*^+/^ vs. *AURKB*^+/−^;*TP53^−^*^/−^ cells, treated with barasertib). P-values from a Chi-squared test with Holm-Sidak’s correction for multiple comparisons. n.s.=non-significant. Data represents pooled results, n=4 biological repeats. **B,** Comparison of the fraction of mitotic failure events between the CAL51 models exposed to 1nM barasertib for 48 hours. Each point represents one biological repeat, calculated as the number of mitotic failures divided by the total mitoses within that replicate. *P*=0.018 (Control vs. *AURKB*^+/−^); *P*=0.018 (Control vs. *TP53^−^*^/−^); *P*=0.016 (*TP53^−^*^/−^ vs. *AURKB*^+/−^;*TP53^−^*^/−^). P-values from repeated-measures one-way ANOVA with Holm-Sidak’s correction for multiple comparisons. Data represents pooled results, n=4 biological repeats. **C,** Representative images of a flow cytometry-based cell cycle analysis of the CAL51 models exposed to 10nM barasertib for 48 hours. WGD negative and positive cells are shown; arrows denote ploidy. **D,** Quantification of the % WGD-positive cells with DNA content greater than 4N. Heterozygous *AURKB*-KO sensitized *TP53-*WT cells to barasertib, increasing the prevalence of WGD events compared to *AURKB*-WT cells. *TP53*-null cells were more resistant to the drug, but heterozygous *AURKB*-KO re-sensitized *TP53-*null cells to the drug. *P*=n.s. (Control vs. *AURKB*^+/−^); *P*<0.0001 (Control vs. *TP53^−^*^/−^); *P*=0.0001 (*TP53^−^*^/−^ vs. *AURKB*^+/−^;*TP53^−^*^/−^). P-values from repeated-measures one-way ANOVA with Holm-Sidak’s correction for multiple comparisons; n=9 biological repeats. n.s.=non-significant. **E,** Quantification of the % of Annexin-positive cells treated with either DMSO (control) of 10nM barasertib. The Δ Annexin-positive cell percentage was calculated by subtracting the percentage in DMSO-treated samples from that in barasertib-treated samples. *P*=0.027 (Control vs. *AURKB*^+/−^); *P*=0.011 (Control vs. *TP53^−^*^/−^); *P*=0.251 (*TP53^−^*^/−^ vs. *AURKB*^+/−^;*TP53^−^*^/−^). P-values from repeated-measures one-way ANOVA with Holm-Sidak’s correction for multiple comparisons; n=7 biological repeats. **F,** Schematic representation of *TP53* inactivation and its effects on AURKB expression and sensitivity to AURKBi. When *TP53* inactivation occurs through point mutations, AURKB expression levels are increased, resulting in increased resistance to AURKBi (top). However, when *TP53* inactivation occurs through Del17p, AURKB expression levels are reduced, leading to increased sensitivity of the cells to AURKBi.

These results indicate that AURKB dependency in Del17p tumors persists even in the context of complete *TP*53 LOF, which is very common in breast cancer. Thus, targeting AURKB may represent a viable therapeutic approach for Del17p mammary tumors, even on the background of a *TP53* mutation. Furthermore, whereas p53 inactivation was previously suggested to increase the resistance of cancer cells to AURKBi, our results reveal that *the mode* of p53 inactivation must be considered; if the biallelic inactivation of p53 involves Del17p, then inhibiting AURKB may remain a viable therapeutic option (**Fig. 8F**).

## Discussion

Our study demonstrates that Del17p is a frequent and clinically meaningful aneuploidy in breast cancer, which is strongly associated with poor prognosis, reduced p53 activity, and increased proliferation. Importantly, Del17p tumors exhibit a specific vulnerability to AURKBi due to AURKB hemizygosity, highlighting AURKB as a potential therapeutic target in Del17p breast cancer. These findings suggest that Del17p status could serve as a predictive biomarker for identifying patients who may benefit the most from AURKB-targeted therapies.

Furthermore, in contrast to many other oncogenes, which become more essential (**39-41**) upon overexpression, we reveal that AURKB behaves similarly to a CYCLOPS gene, becoming more essential when one of its alleles is lost. Importantly, p53 inactivation *per se* results in increased AURKB expression and increased resistance to AURKBi, and p53 inactivation was therefore previously proposed as a biomarker for AURKBi resistance. Our data strongly suggest that the mode of p53 inactivation matters in this regard – Del17p tumors may still be sensitive to AURKBi despite p53 inactivation. Our study therefore calls for clinical exploration of the efficacy of AURKB inhibitors, such as barasertib, in treating Del17p breast cancer patients.

## Supplementary Data

**Supplementary Figure S1: Prevalence of Del17p in breast cancer and its association with tumor features, related to Figure 1**.

**A,** Prevalence of Del17p across breast cancer molecular subtypes in the METABRIC dataset, showing it is a common event across molecular subtypes. **B,** Comparison of the overall survival between patients with Del17p tumors and patients with WT17p tumors, stratified by molecular subtype. Del17p is associated with worse prognosis across the molecular subtypes. P-values from a Log-rank test. **C,** Prevalence of Del17p across breast cancer tumors from the TCGA dataset. **D,** Comparison of disease-free survival between patients with Del17p tumors and patients with WT17p tumors, using the TCGA dataset (n=768). *P*=0.17 from a Log-rank test. **E,** Comparison of the tumor T-stage between Del17p tumors and WT17p tumors, using the TCGA dataset (n=764). *P*<0.0001 from a Chi-squared test.

**Supplementary Figure S2: Del17p is associated with p53 pathway downregulation and increased tumor aggressiveness, related to Figure 2**.

**A,** Prevalence of Del17p and *TP53* mutations (see Material and Methods) in human primary breast cancer samples from the METABRIC dataset, stratified by their molecular subtypes. **B,** Comparison of overall survival between METABRIC patients stratified by their Del17p and *TP53* mutation status, within each molecular subtype, showing worse survival in the patients with Del17p;*TP53*-WT tumors in comparison to those with WT17p;*TP53*-WT tumors, and the worst survival in the patients with Del17p;*TP53*-mut tumors. P-values from a Log-rank test. **C,** Prevalence of Del17p and *TP53* mutation in breast cancer tumors from the TCGA dataset. **D,** Comparison of disease-free survival between TCGA patients stratified by their Del17p and *TP53* mutation status, showing worse survival in the patients with Del17p;*TP53*-WT tumors in comparison to those with WT17p;*TP53*-WT tumors (*P*=0.2248), and the worst survival in the patients with Del17p;*TP53*-mut tumors (*P*=0.4481 when compared to the Del17p;*TP53*-WT tumors). n=667 tumors, P-values from a Log-rank test. **E**, Comparison of the tumor T-stage between TCGA patients stratified by their Del17p and *TP53* mutation status, showing a higher tumor T-stage in the Del17p;*TP53*-WT tumors compared to WT17p;*TP53*-WT and the highest tumor T-stage in the Del17p;*TP53-*mut tumors. n=757 tumors, *P*<0.0001 from a Chi-squared test.

**Supplementary Figure S3: Del17p is associated with upregulation of signatures associated with tumor proliferation, related to Figure 2**.

**A,** GSEA of the METABRIC dataset, showing enrichment for cell cycle-related pathways (e.g., DNA replication, cell cycle checkpoints) in tumors with: Del17p compared to tumors with WT17p (left); Del17p;*TP53*-WT compared to tumors with WT17p;*TP53*-WT (middle); Del17p;*TP53*-mut compared to tumors with Del17p;*TP53*-WT (right). Plot shows enrichment for the ‘Reactome’ gene set. **B,** GSEA of the TCGA dataset, showing enrichment for cell cycle-related pathways (e.g., G2M checkpoint, E2F targets) in tumors with: Del17p compared to tumors with WT17p (left); Del17p;*TP53*-WT compared to tumors with WT17p;*TP53*-WT (middle); Del17p;*TP53*-mut compared to tumors with Del17p;*TP53*-WT (right). Plot shows enrichment for the ‘Hallmark’ gene sets. **C,** GSEA of the TCGA dataset, showing enrichment for cell cycle-related pathways (e.g., DNA replication, cell cycle checkpoints) in tumors with: Del17p compared to tumors with WT17p (left); Del17p;*TP53*-WT compared to tumors with WT17p;*TP53*-WT (middle); Del17p;*TP53*-mut compared to tumors with Del17p;*TP53*-WT (right). Plot shows enrichment for the ‘Reactome’ gene sets.

**Supplementary Figure S4: Del17p is associated with downregulation of genes that reside on Chr17p, related to Figure 2**.

**A,** GSEA of the METABRIC dataset, showing strong depletion for genes residing on Chr17p when this chromosome-arm is lost, both when comparing all tumors (left) and when considering only *TP53*-WT tumors (right). NES=-2.98, q<0.0001; NES=-3.18, q<0.0001, respectively. **B,** GSEA of the TCGA dataset, showing strong depletion for genes residing on Chr17p when this chromosome-arm is lost, both when comparing all tumors (left) and when considering only *TP53*-WT tumors (right). NES=-3.19, q<0.0001; NES=-3.2, q<0.0001.

**Supplementary Figure S5: Reduced expression of AURKB in Del17p breast cancer cell lines, related to Figure 3**.

**A,** Comparison of AURKB mRNA levels between WT17p and Del17p human breast cancer cell lines. Top: non-controlled (left), *P*=0.0002; controlled for *TP53* mutation status (middle), *P*=0.0003; controlled for Chr17q copy number status (right), P=0.0011. Bottom: controlled for aneuploidy score (AS) (**26**) (left), *P*=0.0002; controlled for doubling time (DT) (**27**) (middle), *P*<0.0001; controlled for breast cancer molecular subtype (right), *P*=0.0008. P-values from a two-tailed unpaired Student’s t-test (when controlled for 17q and DT) or from a two-tailed Mann-Whitney U-test (for all other comparisons). **B,** Comparison of AURKB protein level between WT17p and Del17p breast cancer cell lines. Top: non-controlled (left), *P*=0.0334; controlled for *TP53* mutation status (middle), *P*=0.031; controlled for Chr17q copy number status (right), *P*=0.0027. Bottom: controlled for aneuploidy score (AS) (**26**) (left), *P*=0.0334; controlled for doubling time (DT) (**27**) (middle), *P*=0.0364; controlled for breast cancer molecular subtype (right), *P*=n.s.. P-values from a two-tailed Student’s t-test. n.s.=non-significant.

**Supplementary Figure S6: Increased sensitivity of Del17p breast cancer cell lines to RNAi-mediated knockdown of AURKB, related to Figure 3**.

**A,** Comparison of sensitivity to RNAi-mediated knockdown of AURKB between WT17p and Del17p human breast cancer cell lines, based on the essentiality (DEMETER) scores from the RNAi-Novartis screen (**27**). Top: non-controlled (left), *P*=0.0496; controlled for *TP53* mutation status (middle), *P*=0.0029; controlled for Chr17q copy number status (right), *P*=0.0019. Bottom: controlled for aneuploidy score (AS) (**26**) (left), *P*=0.0035; controlled for doubling time (DT) (**27**) (middle), *P*=0.0042; controlled for breast cancer molecular subtype (right), *P*=0.0048. P-values from a two-tailed Mann-Whitney U-test (when non-controlled and controlled for breast cancer molecular subtype) or from a two-tailed Student’s t-test (all other comparisons). **B,** Comparison of sensitivity to RNAi-mediated knockdown of AURKB between WT17p and Del17p breast cancer cell lines, based on the essentiality (DEMETER) scores from the RNAi-Broad screen (**27**). Top: non-controlled (left), *P*=0.0097; controlled for *TP53* mutation status (middle), *P*=0.0367; controlled for Chr17q copy number status (right), *P*=0.0339. Bottom: controlled for aneuploidy score (AS) (**26**) (left), *P*=0.0075; controlled for doubling time (DT) (**27**) (middle), *P*=0.0119; controlled for breast cancer molecular subtype (right), *P*=0.0362. P-values from a two-tailed Mann-Whitney U-test (when controlled for AS) or from a two-tailed unpaired Student’s t-test (all other comparisons). **C,** Comparison of sensitivity to RNAi-mediated knockdown of AURKB between WT17p and Del17p breast cancer cell lines, based on the essentiality (DEMETER) scores from the RNAi-combined results (**42**). Top: non-controlled (left), *P*=0.025; controlled for *TP53* mutation status (middle), *P*=0.0395; controlled for Chr17q copy number status (right), *P*=0.0365. Bottom: controlled for aneuploidy score (AS) (**26**) (left), *P*=0.0091; controlled for doubling time (DT) (**27**) (middle), *P*=0.0003; controlled for breast cancer molecular subtype (right), *P*=n.s.. P-values from a two-tailed unpaired Student’s t-test. n.s.=non-significant.

**Supplementary Figure S7: Increased sensitivity of Del17p breast cancer cell lines to CRISPR-mediated knockout of *AURKB,* related to Figure 3**.

**A,** Comparison of sensitivity to CRISPR-mediated knockout of AURKB between WT17p and Del17p human breast cancer cell lines, based on the essentiality (CERES) scores from the CRISPR-Avana screen. Top: non-controlled (left), *P*=0.0146; controlled for *TP53* mutation status (middle), *P*=0.0145; controlled for Chr17q copy number status (right); *P*=0.0075. Bottom: controlled for aneuploidy score (AS) (**26**) (left), *P*=0.013; controlled for doubling time (DT) (**27**) (middle), *P*=n.s.; controlled for breast cancer molecular subtype (right), *P*=0.002. P-values from a two-tailed unpaired Student’s t-test. **B,** Comparison of sensitivity to CRISPR-mediated knockout of AURKB between WT17p and Del17p breast cancer cell lines, based on the essentiality (CERES) scores from the CRISPR-Sanger screen. Top: non-controlled (left), *P*=n.s.; controlled for *TP53* mutation status (middle), *P*=n.s.; controlled for Chr17q copy number status (right), *P*=n.s.; Bottom: controlled for aneuploidy score (AS) (**26**) (left), *P*=n.s.; controlled for doubling time (DT) (**27**) (middle), *P*=n.s.; controlled for breast cancer molecular subtype (right), *P*=n.s.. P-values from a two-tailed unpaired Student’s t-test. n.s.=non-significant.

**Supplementary Figure S8: Comparison of the sensitivity of MD-AMB-231 and MD-AMB-468 to multiple cancer-related drugs, related to Figure 4**.

**A,** Immunofluorescence images of MD-AMB-231 (left) and MD-AMB-468 (right) cells, showing lower AURKB protein expression levels (red) in the nuclei of MD-AMB-468 interphase cells. DAPI (blue). **B,** Quantification of (A). *P*=0.0064 from a one-sample t-test. **C,** Comparison of the sensitivity of MD-AMB-231 and MD-AMB-468 cells to a battery of cancer-related drugs. Although MD-AMB-468 was more sensitive to most drugs, barasertib displayed the largest difference. Top: Cycloheximide (left), Doxorubicin (right). Bottom: Paclitaxel (left), Etoposide (right). Cells were exposed to the drugs for 72 hours. Each curve shows the average of 3 (Cycloheximide, Paclitaxel) or 4 (Doxorubicin, Etoposide) biological repeats. **D,** Ratios of the Area under the curve (AUC) values between MD-AMB-468 and MD-AMB-231 for each of the drugs tested in (C). *P*=n.s. (Barasertib vs. Doxorubicin; Baraserib vs. Etoposide; Barasertib vs. Paclitaxel); *P*=0.013 (Barasertib vs. Cycloheximide). P-values from a one-way Kruskal-Wallis test with Dunn’s correction for multiple comparisons. n=3 (Cycloheximide, Paclitaxel) or 4 (Doxorubicin, Etoposide) biological repeats. n.s.=non-significant.

**Supplementary Figure S9: Del17p increases sensitivity to α-amanitin, YM-155 and barasertib in MCF7 cells, related to Figure 5**.

**A,** Comparison of AURKB mRNA levels across the MCF7 strains using RT-qPCR. *P*=0.0026 (N vs. D) or *P*=n.s. (all other comparisons). P-values from a one-sample t-test, n=5 biological repeats. **B,** Comparison of AURKB protein levels across the MCF7 strains using WB. *P*=n.s. (E vs. D); *P*=0.0277 (K vs. D); *P*=0.036 (L vs. D); *P*=0.0055 (O vs. D); *P*=n.s. (N vs. D). P-values from a one-sample t-test. n=4 biological repeats. n.s.=non-significant. **C,** Dose-response curves comparing the response of WT17p and Del17p strains to α-amanitin. Left: Individual curves for each MCF7 strain. Right: Average curves for WT17p and Del17p strains. n=3 biological repeats. **D,** Comparison of the Area Under the Curve (AUC) values from the experiments shown in (C). *P*=0.05 from a one-tailed Mann-Whitney U-test. n=3 biological repeats. **E,** Volcano plot comparing the sensitivity of WT17p strains and Del17p strains to 27 drugs. Sensitivity data were obtained from a drug screen against multiple MCF7 strains (**32**). The x-axis represents the magnitude of the effect difference, the y-axis represents the statistical significance. The most significantly-differential drug is the Survivin inhibitor YM-155 (red), to which Del17p strains are significantly more sensitive to. **F,** Dose-response curves comparing the response of WT17p and Del17p strains to YM-155. Data obtained from the MCF7 drug screen (**32**). **G,** Dose-response curves comparing the response of WT17p and Del17p strains to YM-155. Left: Individual curves for each MCF7 strain. Right: Average curves for WT17p and Del17p strains. n=3 biological repeats. **H,** Comparison of the Area Under the Curve (AUC) values from the experiments shown in (G, Right). *P*=0.0194 from a one-tailed unpaired Student’s t-test. n=3 biological repeats. **I,** Dose-response curves comparing the response of WT17p and Del17p MCF7 strains exposed to 10nM barasertib for 72 hours. Data represents pooled results, n=3 biological repeats.

**Supplementary Figure S10: Generation of heterozygous *AURKB*-KO breast cancer cell lines, related to Figure 7 and Figure 8**.

**A,** Fraction of CAL51 single-cell clones with a fusion band indicative of a loss of exon 2. No homozygous AURKB-KO clones have been recovered. **B,** Comparison of AURKB mRNA levels between *lacZ*-control (Control) and three heterozygous *AURKB*-KO CAL51 clones (*AURKB*^+/−^), using RT-qPCR. **C,** Comparison of AURKB protein levels between Control and three heterozygous *AURKB*-KO CAL51 clones, using WB. α-tubulin used as a control. **D,** Quantification of the WB results shown in (C). **E,** Immunofluorescence images of Control and heterozygous *AURKB*-KO CAL51 cells, showing reduced protein expression of AURKB in the nuclei of the heterozygous *AURKB*-KO cells. **F,** Quantification of the immunofluorescence shown in (E). *P*<0.0001 (Control vs. 10); *P*<0.0001 (Control vs. 24); *P*<0.0001 (Control vs. 25). P-values from a one-way Kruskal-Wallis test with Dunn’s correction for multiple comparisons. **G,** Sanger sequencing of the fusion (left) and WT (right) PCR products from CAL51 single-cell clones. Left: Primers flanking the expected fusion junction between introns 1 and 2 were used, demonstrating the fusion of the sequence proximal to the first cut site with the sequence distal to the second cut site. Right: Primers flanking exon 2, which was not excised on the other, intact allele. **H,** Comparisons of *TP53* (left) and CDKN1A (right) mRNA expression levels in Control CAL51 cells or in cells with heterozygous *AURKB*-KO, on a *TP53*-WT or *TP53*-null background, demonstrating successful *TP53* knockout in the *TP53*-null clones. Left: *P*=0.0232 (Control vs. *TP53^−^*^/−^); *P*=0.0046 (*AURKB*^+/−^ vs. *AURKB*^+/−^;*TP53^−^*^/−^). Right: *P*=0.0033 (Control vs. *TP53^−^*^/−^ and *AURKB*^+/−^ vs. *AURKB*^+/−^;*TP53^−^*^/−^). P-values from repeated-measures one-way ANOVA with Holm-Sidak’s correction for multiple comparisons. n=3 biological repeats. **I,** Comparisons of p53 and p21 protein levels between Control and heterozygous *AURKB*-KO CAL51 cells, on a *TP53*-WT or *TP53*-null background, using WB. α-tubulin used as a control. The results demonstrate successful *TP53* knockout in the *TP53*-null clones.

**Supplementary Figure S11: *AURKB* hemizygosity confers sensitivity to AURKB and Survivin inhibitors in a p53-dependent manner, related to Figure 7**.

**A,** Dose-response curves comparing the response to YM-155 of Control and heterozygous *AURKB*-KO CAL51 cells, on a *TP53*-WT or *TP53*-null background. *TP53*-null cells are more resistant to the drug, but AURKB hemizygosity increases the sensitivity on both *TP53* backgrounds. Data represents pooled results, n=6 biological repeats. **B,** Comparison of the Area Under the Curve (AUC) values from the experiments shown in (J). *P*=0.0001 (Control vs. *AURKB*^+/−^); *P*=0.0271 (Control vs. *TP53^−^*^/−^); *P*<0.0001 (*AURKB*^+/−^ vs. *AURKB*^+/−^;*TP53^−^*^/−^). P-values from repeated-measures one-way ANOVA with Holm-Sidak’s correction for multiple comparisons; n=6 biological repeats.

**Supplementary Figure S12: Gating strategy for flow cytometry analyses, related to Figure 8**.

**A,** CAL51 heterozygous *AURKB*-KO cells treated with DMSO (control; top panels) or 10nM barasertib (bottom panels), followed by Hoechst staining for DNA content determination. The singlets gate was determined using the barasertib-treated cells (bottom left) and applied to the control cells (top left). For ploidy gating, a strategy based on hepatocyte gating (**43**) was used (right panels). Cells with DNA content greater than 4N (>4N) were classified as WGD-positive, while cells with DNA content between 2N and 4N (2N-4N) were classified as WGD-negative. The singlets gate was common across all cell lines, whereas the WGD-positive gate was determined individually for each cell line. **E,** Representative flow cytometry plots showing Annexin-positive cells treated with either DMSO (control) or 10nM barasertib. For each cell line, autofluorescence was assessed using unstained samples (left), and apoptosis was measured using Annexin staining (right). The percentage of apoptotic cells for each condition was calculated by subtracting the signal of the unstained sample from the stained sample. Shown are *TP53*-null cells.

**Supplementary Table S1**

Antibodies, drugs, primers, CRISPR guides and plasmids used in this study.

**Supplementary Table S2**

Summary of Gene Set Enrichment Analysis (GSEA) results.

**Supplementary Table S3**

DepMap two-group comparisons across multiple expression and dependency datasets.

## Data Availability

The TCGA and METABRIC data were obtained from cBioPortal (https://www.cbioportal.org/study/summary?id=brca_tcga_pan_can_atlas_2018; https://www.cbioportal.org/study/summary?id=brca_metabric). Datasets used: ‘TCGA: *data_mrna_seq_v2_rsem.csv’*; METABRIC: ‘*data_mrna_illumina_microarray.csv’*. DepMap data were obtained from https://depmap.org/portal/. For the DepMap analysis, the following version 20Q3 files were used: ‘*Achilles_gene_effect.csv*, *CCLE_expression.csv’*, ‘*CCLE_mutations.csv’*, ‘*CCLE_segment_cn.csv’*, ‘*Protein_quant_current_normalized.csv’*, ‘*Gene_effect*.csv’, ‘*D2_DRIVE_gene_dep_scores.csv’*, ‘*D2_Achilles_gene_dep_scores.csv’* and ‘*D2_combined_gene_dep_scores.csv’*. All datasets are available within the article and its Supplementary Tables or from the corresponding author upon request. The code used for data analysis is available at: https://github.com/BenDavidLab/17p_del_BRCA/tree/main.

## Authors’ Contributions

U.B.-D. conceived the study. T.W. led the computational analyses. T.W. and E.S. performed the *in vitro* experiments. R.S., G.W., K.L. and G.L. assisted with the computational analyses. H.O. and K.L. assisted with the *in vitro* experiments. T.W., E.S. and U.B.-D. wrote the manuscript with inputs from all authors. U.B.-D., T.W. and E.S. designed the figures and E.S. prepared the figures. U.B.-D. Supervised the study.

## Acknowledgments

We thank the members of the Ben-David laboratory for fruitful discussions. Work in the Ben-David laboratory is supported by the Israel Cancer Research Fund Project Award, the Israel Science Foundation project grant (1805/21), the BSF project grant (2019228) and the European Research Council Starting Grant (945674). E.S. is supported by the Eugenia Gelman Estate for Cancer Research PhD Fellowship, H.O. is supported by the JSPS Overseas Research Fellowship, and R.S is supported by the Edmond J. Safra Center for Bioinformatics at Tel Aviv University.

